# Lineage specific histories of *Mycobacterium tuberculosis* dispersal in Africa and Eurasia

**DOI:** 10.1101/210161

**Keywords:** phylogeography, evolution, pathogen, migration, demography

## Abstract

*Mycobacterium tuberculosis* (*M.tb*) is a globally distributed, obligate pathogen of humans that can be divided into seven clearly defined lineages. Identifying how the ancestral clone of *M.tb* spread and differentiated is important for identifying the ecological drivers of the current pandemic. We reconstructed *M.tb* migration in Africa and Eurasia, and investigated lineage specific patterns of spread. Applying evolutionary rates inferred with ancient *M.tb* genome calibration, we link *M.tb* dispersal to historical phenomena that altered patterns of connectivity throughout Africa and Eurasia: trans-Indian Ocean trade in spices and other goods, the Silk Road and its predecessors, the expansion of the Roman Empire and, the European Age of Exploration. We find that Eastern Africa and Southeast Asia have been critical in the dispersal of *M.tb*. Our results reveal complex relationships between spatial dispersal and expansion of *M.tb* populations, and delineate the independent evolutionary trajectories of bacterial sub-populations underlying the current pandemic.

## Introduction

The history of tuberculosis (TB) has been rewritten several times as genetic data accumulate from its causative agent, *Mycobacterium tuberculosis* (*M.tb*). In the nascent genomic era, these data refuted the long-held hypothesis that human-adapted *M.tb* emerged from an animal adapted genetic background represented among extant bacteria by *Mycobacterium bovis*, another member of the *Mycobacterium tuberculosis* complex (MTBC) (Brosch et al. 2002). Genetic data from bacteria infecting multiple species of hosts revealed that currently known non-primate-adapted strains form a nested clade within the diversity of extant *M.tb* (Behr et al. 1999; Brosch et al. 2002; Hershberg et al. 2008).

*M.tb* can be classified into seven well-differentiated lineages, which differ in their geographic distribution and association with human sub-populations (Hirsh et al. 2004; Gagneux et al. 2006). This observation led to the hypothesis that *M.tb* diversity has been shaped by human migrations out of Africa, and that the most recent common ancestor (MRCA) of extant *M.tb* emerged in Africa approximately 73,000 years ago, coincident with estimated waves of human migration (Comas et al. 2013). Human out of Africa migrations are a plausible means by which *M.tb* could have spread globally. However, *M.tb* evolutionary rate estimates based on a variety of calibration methods are inconsistent with the out of Africa hypothesis (Eldholm et al. 2016; Brynildsrud et al. 2018).

When calibrated with ancient DNA, the estimates of the time to most recent common ancestor (TMRCA) for the MTBC are <6,000 years before present (Bos et al. 2014; Kay et al. 2015). This is not necessarily the time period over which TB first emerged, as it is possible – particularly given the apparent absence of recombination among *M.tb* (Pepperell et al. 2013) – that the global population has undergone clonal replacement events that displaced ancient diversity from the species.

*M.tb* is an obligate pathogen of humans with a global geographic range. The finding of a recent origin for the extant *M.tb* population raises the question of how the organism could have spread within this timeframe to occupy its current distribution. *M.tb* populations in the Americas show the impacts of European colonial movements as well as recent immigration (Pepperell et al. 2011; Brynildsrud et al. 2018); the role of other historical phenomena in driving TB dispersal is not well understood. Here we sought to reconstruct the migratory history of *M.tb* populations in Africa and Eurasia within the framework of a recent origin and evolutionary rates derived from ancient DNA data (Bos et al. 2014; Kay et al. 2015). We discovered lineage-specific patterns of migration and a complex relationship between *M.tb* effective population growth and migration. Our results connect *M.tb* migration to major historical events in human history that altered patterns of connectivity in Africa and Eurasia. These findings provide context for a recent evolutionary origin of the MRCA of *M.tb* (Pepperell et al. 2013; Bos et al. 2014; Kay et al. 2015), which represents yet another paradigm shift in our understanding of the history and origin of this successful pathogen.

## Results

### Genetic and geographic structures of global M.tb populations

In order to establish the contemporary geographic distributions of *M.tb* lineages, we translated the spoligotypes reported for 42,358 *M.tb* isolates to their corresponding lineage designations (fig. 1). Geographic patterns in prevalence vary between lineages. Lineage 1 (L1) is prevalent in regions bordering the Indian Ocean, extending from Eastern Africa to Melanesia. Lineage 2 (L2) is broadly distributed, with a predominance in Eastern Eurasia and South East Asia.

**Fig. 1.**
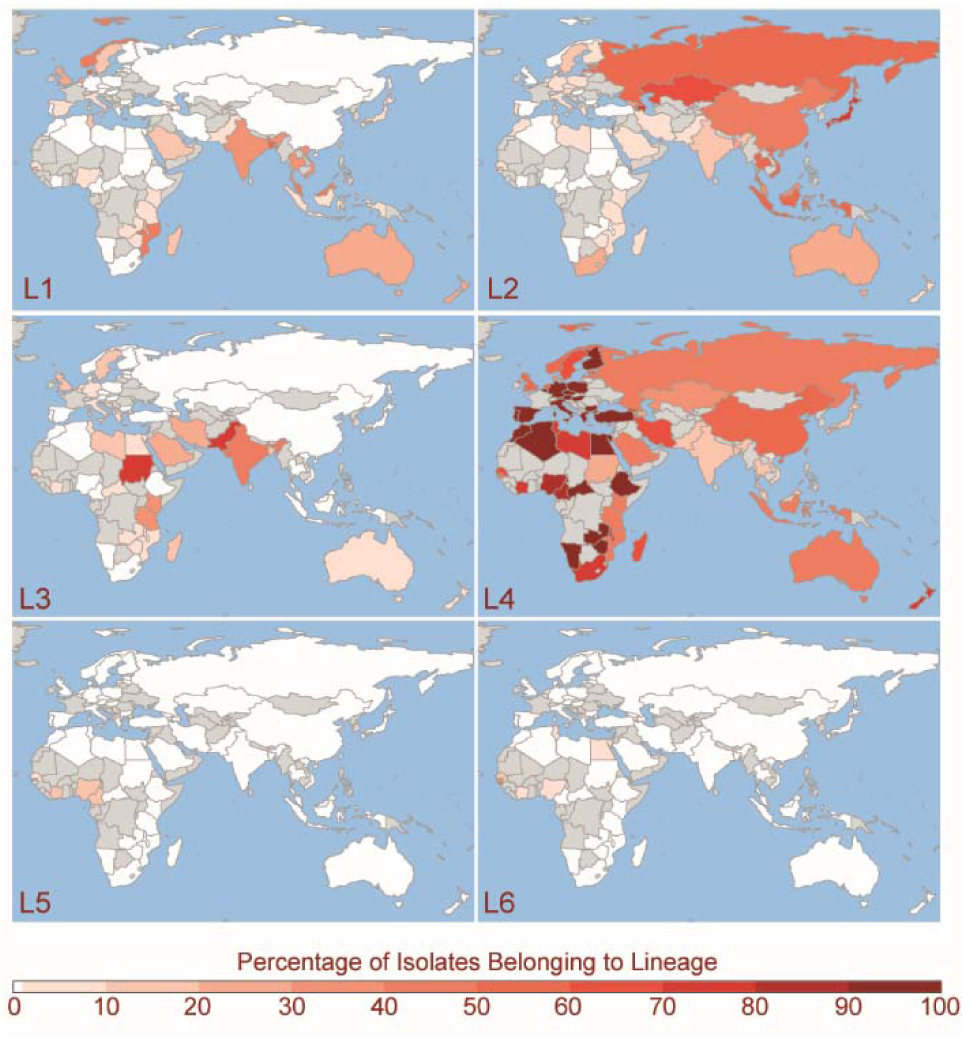
Geographic distributions of *Mycobacterium tuberculosis* lineages 1-6. Spoligotypes from the SITVIT WEB database (*n* = 42,358) were assigned to lineages 1-6. Countries are colored from white to dark-red based on the percentage of isolates from the country belonging to each lineage. Unsampled countries and those with less than 10 isolates in the database are shown in grey. Lineage 7 (not pictured) is found exclusively in Ethiopia.

Lineage 3 (L3) is similar to L1 in that its distribution rings the Indian Ocean, but it does not extend into Southeastern Asia, it has a stronger presence in Northern Africa, and a broader distribution across Southern Asia. Lineage 4 (L4) is strikingly well dispersed, with a predominance throughout Africa and Europe and the entire region bordering the Mediterranean. Lineages 5 (L5) and 6 (L6) are found at low frequencies in Western and Northern Africa. Lineage 7 (L7), as previously described (Blouin et al. 2012; Firdessa et al. 2013; Comas et al. 2015), is limited to Ethiopia.

We compiled a diverse collection of *M.tb* genomes for phylogenetic and population genetic inference of the demographic and migratory history of the extant *M.tb* population (*see Methods*). Our dataset consists of whole-genome sequences (WGS) from 552 *M.tb* isolates collected from 51 countries (spanning 13 UN geoscheme subregions), which we refer to as the Old World collection (fig. S1, table S1). We included sites in the alignment where at least half of these isolates had confident data (60,787 variant sites; 3,838,249 bp) for subsequent analyses, unless otherwise noted.

The inferred maximum likelihood phylogeny reveals the well described *M.tb* lineage structure, and some associations are evident between lineages and geographic regions (defined here by the United Nations geoscheme) (fig. S2). The phylogeny has an unbalanced shape, with long internal branches that define the lineages and feathery tips, suggestive of recent population expansion.

Genetic diversity, as measured by the numbers of segregating sites and pairwise differences (Watterson’s C and π), varied among lineages (table 1). L1 and L4 group together and have the highest diversity; L2, L3, L5, and L6 have similar levels of diversity and form the middle grouping; L7 has the lowest diversity. We used an analysis of molecular variance (AMOVA) to delineate the effects of population sub-division on *M.tb* diversity (table 1). The Old World collection was highly structured among UN subregions (21% of variation attributable to between-region comparisons), whereas this structure was less apparent when regions were defined by the botanical contents outlined by the World geographic scheme for recording plant distributions (14%). This is consistent with *M.tb’*s niche as an obligate human pathogen, with bacterial population structure directly shaped by that of its host population (i.e. reflected in UN subregions) rather than climatic and other environmental features (reflected in botanical continent definitions). We obtained similar results when the lineages were considered separately, except for L4, which had little evidence of population structure (4% variation among UN subregions, 2% among botanical continents).

**Table 1.**
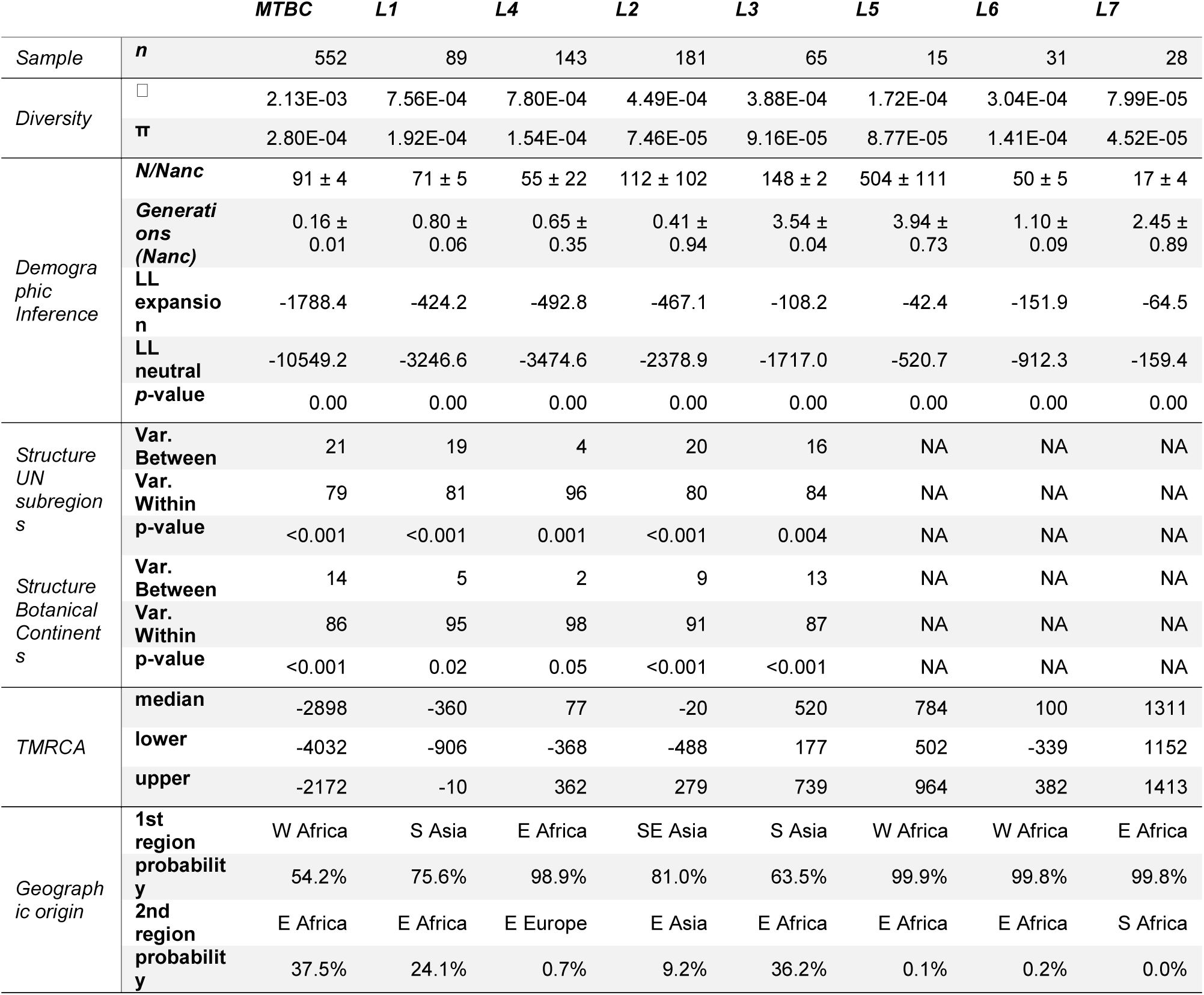
Genetic diversity of Old World *M.tb* across lineages 1-7. TMRCA estimates reflect scaling of results to evolutionary rates calibrated from ancient DNA [median 5.00×10^−8^ substitutions/ site/ year (Kay et al. 2015) and are written as calendar years. To account for uncertainty in this rate estimate, our lower and upper TMRCA estimates reflect scaling of our results with the low and high bounds of the 95% highest posterior density estimates of the rate reported from ancient DNA analysis (i.e. 4.06×10^−8^ and 5.87×10^−8^, respectively).

### Distinct demographic histories of the M.tb lineages

Bayesian inferred trees vary among lineages (fig. 2), likely reflecting their distinct demographic histories. Branch lengths are relatively even across the phylogenies of L1 and L4, whereas L2 and L3 have a less balanced structure. The long, sparse internal branches and radiating tips of L2 and L3 phylogenies are consistent with an early history during which the effective population size remained small (and diversity was lost to drift), followed by more recent population expansion. L5 has a star-like structure, consistent with rapid population expansion. Jointly inferred Bayesian skyline plot (BSP) reconstructions of effective population sizes over time suggest that lineages 1-6 have undergone expansion (fig. 3 – top panel, fig. S3). We estimate that L2 and L3 underwent abrupt expansion at approximately the same time, whereas expansions of L1 and L4 appeared relatively smooth.

**Fig. 2.**
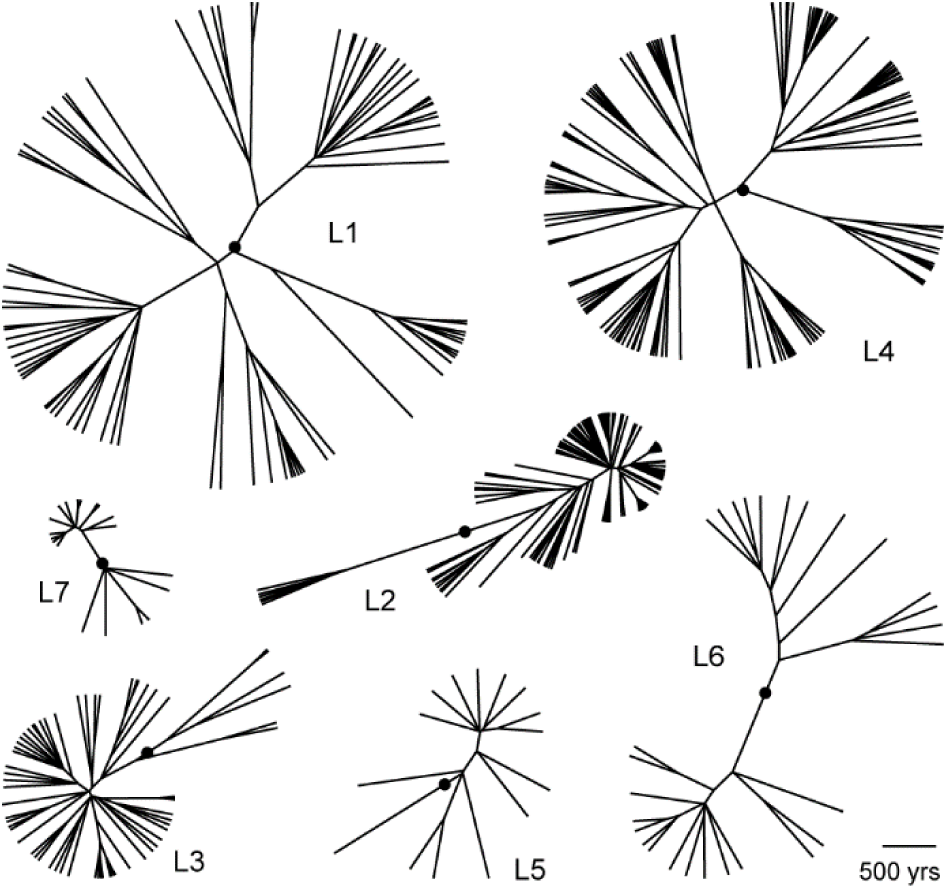
Maximum clade credibility phylogenies of *Mycobacterium tuberculosis* lineages 1-6. Bayesian analyses were performed on each lineage alignment with the general time reversible model of nucleotide substitution with a gamma distribution to account for rate heterogeneity between sites, a strict molecular clock, and Bayesian skyline plot demographic models. The most recent common ancestor (MRCA) of each lineage is indicated with a black circle; the MRCA of individual lineage phylogenies were informed by the phylogeny of the entire Old World collection, which was dated using a substitution rate of 5 × 10^−8^ substitutions/site/year (Kay et al. 2015).

**Fig. 3.**
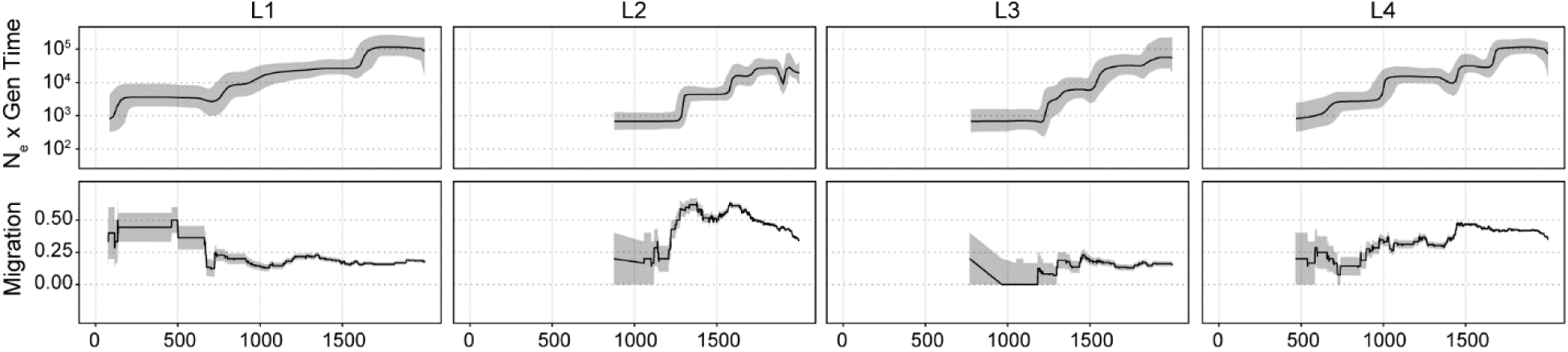
Patterns of effective population size and migration through time of *Mycobacterium tuberculosis* lineages 1-4. Bayesian skyline plots (top panels) show inferred changes in effective population size (N_e_) through time deduced from lineage specific analyses. Black lines denote median N_e_ and gray shading the 95% highest posterior density. Estimated migration through time (see *Methods*) for each lineage is shown in the bottom panels (see *Methods*). Grey shading depicts the rates inferred after the addition or subtraction of a single migration event, and demonstrate the uncertainty of rate estimates, particularly from the early history of each lineage. Dates are shown in calendar years and are based on scaling the phylogeny of the Old World collection with a substitution rate of 5 × 10^−8^ substitutions/site/year (Kay et al. 2015).

We used the methods implemented in ∂a∂i to reconstruct the demographic histories of each *M.tb* population (i.e. lineage) from its synonymous site frequency spectrum (SFS). As demographic inference with ∂a∂i is sensitive to missing data, loci at which any sequence in the individual lineage alignments had a gap or unknown character were removed for these analyses. Consistent with the BSP analyses performed in BEAST, instantaneous expansion and exponential growth models offered an improved fit to the data in comparison with the constant population size model for each lineage and the entire Old World collection (fig. S4). Parameter estimates varied widely across runs for the exponential growth model, so we report results only for the instantaneous expansion model (table 1).

### Major events in M.tb’s migratory history

There was evidence of isolation by distance in the global *M.tb* population, as assessed with a Mantel test of correlations between genetic and geographic distances. We defined geographic distances using three schemes: great circle distances, great circle distances through waypoints of human migration as described in (Ramachandran et al. 2005), and distances along historical trade routes. Waypoints are used to make distance estimates more reflective of presumed human migration patterns (i.e., when calculating between-continent distances, it is generally thought that humans did not pass through large bodies of water, and thus a waypoint is used). To allow comparisons between the schemes, values were centered and standardized (*see Methods*).

Values of the Mantel test statistic were similar for great circle distances (r = 0.16) and trade network distances (r = 0.16), with distances through waypoints reflective of human migration patterns having a lower value (r = 0.14, *p* = 0.0001 for all three analyses). In analyses of human genetic data, adjustment of great circle distances with waypoints results in a higher correlation between genetic and geographic distances (Ramachandran et al. 2005). Our Mantel test results therefore do not support a pattern of isolation by distance as expected if out of Africa human migrations were the primary influence on global diversity of extant *M.tb* (Comas et al. 2013).

To further investigate a potential influence of ancient human migration on *M.tb* evolution, we calculated the correlation between *M.tb* genetic diversity (π) within subregions and their average distances from Addis Ababa, a proxy for a possible origin of anatomically modern human expansion out of Africa. Contrary to what is observed for human population diversity (Ramachandran et al. 2005), we did not observe a significant decline in *M.tb* diversity as a function of distance in our Old World collection (adjusted R-squared = -0.1, *p* = 0.88), nor when we included samples from the Americas (adjusted R-squared = 8.9 × 10^−4^, *p* = 0.34, note S1, fig. S5, table S2).

We used the methods implemented in BEAST to reconstruct the migratory history of the entire Old World *M.tb* collection as well as individual lineages within it, modelling geographic origin of isolates (UN subregion or country) as a discrete trait (fig. 4, figs. S6-S10). Using an evolutionary rate calibrated with 18^th^ century *M.tb* DNA of 5 × 10^−8^ substitutions/site/year (Kay et al. 2015), which is similar to the rate inferred with data from 1,000 year old specimens (Bos et al. 2014), our estimate of the time to most recent common ancestor for extant *M.tb* is between 4032 BCE and 2172 BCE (table 1; date ranges are based on the upper and lower limits of the 95% highest posterior density (HPD) for the rate reported in Kay et al. (2015) which is more conservative than the 95% HPD of our model). We infer an African origin for the MRCA (Eastern or Western subregion, table 1, fig. 4, fig. S6). Shortly after emergence of the common ancestor, we infer a migration of the L1-L2-L3-L4-L7 ancestral lineage from Western to Eastern Africa (we estimate prior to 2683 BCE), with subsequent migrations occurring out of Eastern Africa.

**Fig. 4.**
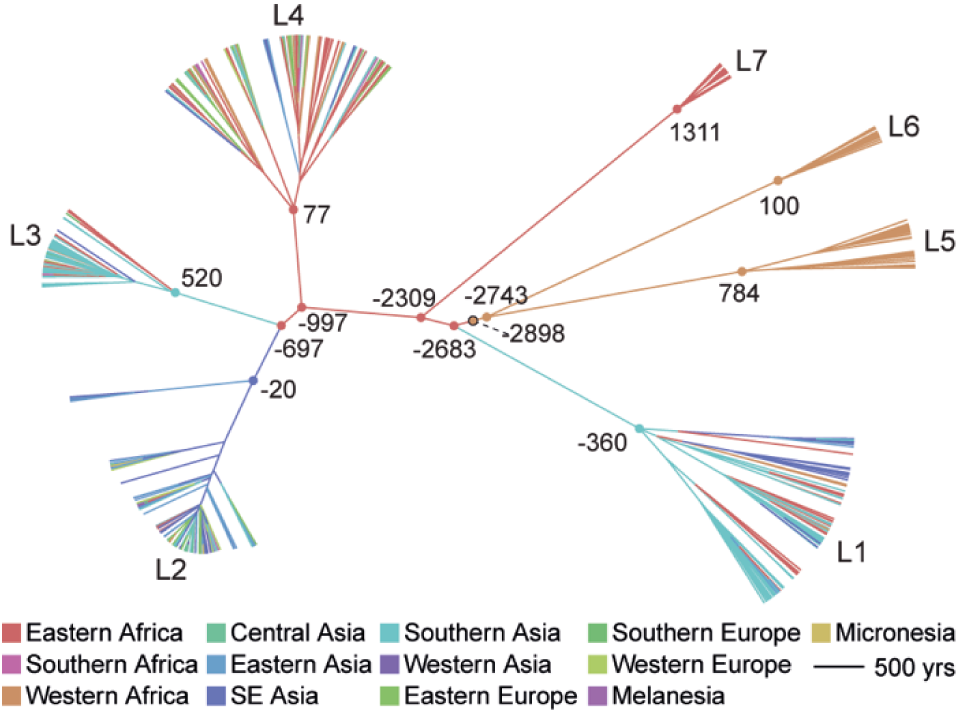
Maximum clade credibility tree of the Old World Collection. Estimated divergence dates are shown in calendar years based on median heights and a substitution rate of 5 × 10^−8^ substitutions/site/year (Kay et al. 2015). Branches are colored according to the inferred most probable geographic origin. Nodes corresponding to the most recent common ancestors (MRCA) of each lineage, lineage splits, and the MRCA of *M. tuberculosis* (outlined black) are marked with circles and colored to reflect their most probable geographic origin.

In our phylogeographic reconstruction, emergence of L1 follows migration from Eastern Africa to Southern Asia at some time between the 3^rd^ millennium and 4^th^ century BCE (table 1, fig. 4, fig. S6). L1 has an ‘out of India’ phylogeographic pattern (fig. S7), with diverse Indian lineages interspersed throughout the phylogeny. This suggests that the current distribution of L1 around the Indian Ocean (fig. 1) arose from migrations out of India, from a pool of bacterial lineages that diversified following migration from Eastern Africa.

The phylogeographic reconstruction further indicates that following the divergence of L1, *M.tb* continued to diversify in Eastern Africa, with emergence of L7 there, followed by L4 (table 1, fig. 4, fig. S6). The contemporary distribution of L4 is extremely broad (fig. 1) and in this analysis of the Old World collection we infer an East African location for the internal branches of L4. Notably, in the lineage-specific analyses, we infer a European location for these branches (fig. S8). The difference is likely due to the fact that inference is informed by deeper as well as descendant nodes in the Old World collection. Together, these results imply close ties between Europe and Africa during the early history of this lineage that we estimate emerged in the 1^st^ century CE (368 BCE-362 CE, table 1).

After the emergence of L1 and L7 from Eastern Africa, our analyses suggest that a migration occurring between 697 BCE and 520 CE established L3 in Southern Asia, with subsequent dispersal out of Southern Asia into its present distribution, which includes Eastern Africa (i.e., a back migration of L3 to Africa, fig. 1). We estimate that L2 diversified in South Eastern Asia following migration from Eastern Africa at some point between 697 BCE and 20 BCE (table 1, fig. 4, fig. S6). Previously published analyses of L2 phylogeography also inferred a Southeast Asian origin for the lineage (Luo et al. 2015; Liu et al. 2018).

### Lineage and region specific patterns of migration

Our phylogeographic reconstruction indicated that temporal trends in migration varied among lineages (fig. 3 – bottom panel). We infer that L1 was characterized by high levels of migration until approximately the 7^th^ century CE, when the rate of migration decreased abruptly and remained stable thereafter. L3, by contrast, exhibited consistently low rates of migration. L2 and L4 had more variable trends in migration, as each underwent punctuated increases in migration rate. Temporal trends in growth and migration are congruent for L2 and L4, with increases in migration rate preceding effective population expansions; this is not the case for L1 and L3. Taken together, these results suggest that L1 and L3 populations (as well as L5 and L6, fig. S3b) grew *in situ*, whereas range expansion may have contributed to the growth of L2 and L4.

We employed the Bayesian stochastic search variable selection method (BSSVS) in BEAST (Lemey et al. 2009) to estimate relative migration rates within the most parsimonious migration matrix. A map showing inferred patterns of connectivity among UN subregions and relative rates of *M.tb* migration with strong posterior support is shown in fig. 5. South Eastern Asia was the most connected region in our analyses, with significant rates of migration connecting it to eight other regions. Eastern Africa, Eastern Europe, and Southern Asia were also highly connected, with significant rates with six, six, and five other regions, respectively. Western Africa, Eastern Asia, and Western Asia were the least connected regions, with just one significant connection each (to Eastern Africa, South Eastern Asia, and Eastern Europe, respectively). Our sample from Western Asia is, however, limited (table S1) and migration from this region may have consequently been underestimated. The highest rates of migration were seen between Eastern Asia and Southeastern Asia, and between Eastern Africa and Southern Asia.

**Fig. 5.**
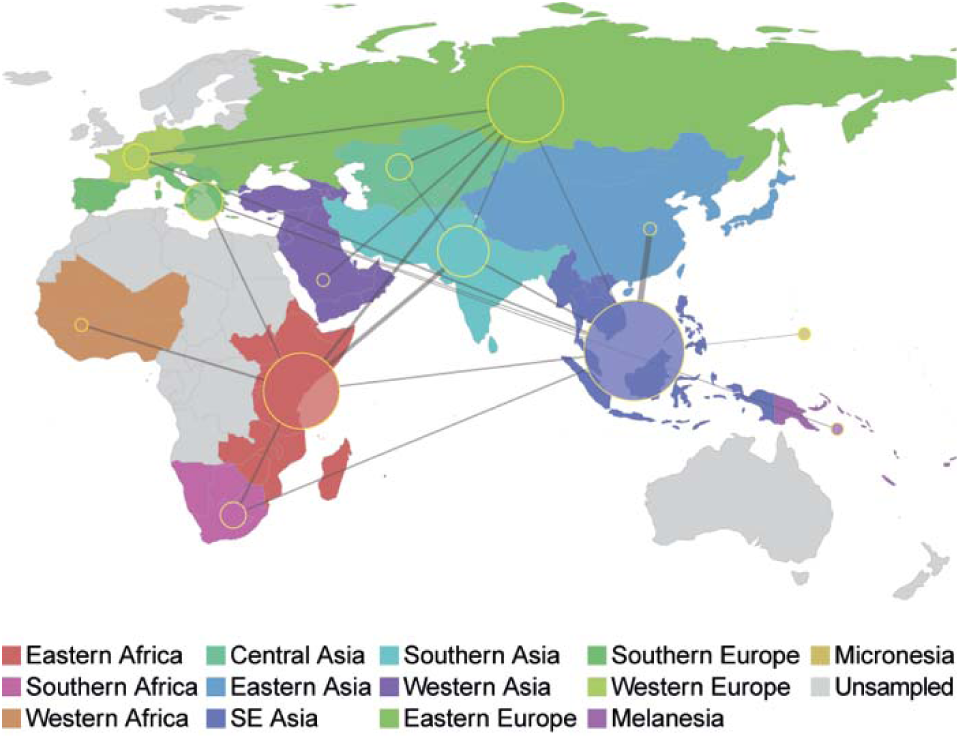
Connectivity of UN subregions during dispersal of *Mycobacterium tuberculosis*. The Bayesian stochastic search variable selection method was used to identify and quantify migrations with strong support in discrete phylogeographic analysis of the Old World collection. Node sizes reflect the number of significant migrations emanating from the region observed in the phylogeny, whereas the thickness of lines connecting regions reflects the estimated relative rate between regions.

Lineage specific analyses suggest that migration between Southern Asia, Eastern Africa, and South Eastern Asia has been important for the dispersal of L1, whereas South Eastern Asia and Eastern Europe have been important for L2 (fig. S11). L3 is similar to L1 in that there is evidence of relatively high rates of migration between Southern Asia and Eastern Africa. There is also evidence of migration within Africa between the eastern and southern subregions. In the analyses of migration for L4, Eastern Africa appeared highly connected with other regions.

## Discussion

Our reconstructions of *M.tb* dispersal throughout the Old World delineate a complex migratory history that varies substantially between bacterial lineages. Patterns of diversity among extant *M.tb* suggest that historical pathogen populations were capable of moving fluidly over vast distances. Using evolutionary rate estimates from ancient DNA calibration, we time the dispersal of *M.tb* to a historical period of exploration, trade, and increased connectivity among regions of the Old World.

Consistent with prior reports (Comas et al. 2013), we infer an origin of *M.tb* on the African continent (table 1, fig. 4, fig. S6). There is a modest preference for Western Africa over Eastern Africa (54% versus 38% inferred probability), likely due to the early branching West African lineages (i.e. *Mycobacterium africanum*, L5 and L6). Larger samples may allow more precise localization of the *M.tb* MRCA, and Northern Africa in particular is under-studied.

We infer L1 to be the first lineage that emerged out of Africa; L1 is currently concentrated in regions bordering the Indian Ocean from Eastern Africa to Melanesia (fig. 1). In our phylogeographic reconstruction, the genesis of this lineage traces to migration from Eastern Africa to Southern Asia at some point between the 3^rd^ millennium and 4^th^ century BCE, with subsequent dispersal occurring out of the Indian subcontinent. Our results suggest that the early history of L1 was characterized by high levels of migration, particularly between Southern Asia and Eastern Africa, and between Southern Asia and South Eastern Asia (fig. 3, fig. S11). The geographic distribution of L1, the timing of its emergence and spread, as well as patterns of connectivity underlying its dispersal, are all consistent with migration via established trans-Indian Ocean trade routes linking Eastern Africa to Southern and South Eastern Asia (fig. 6).

**Fig. 6.**
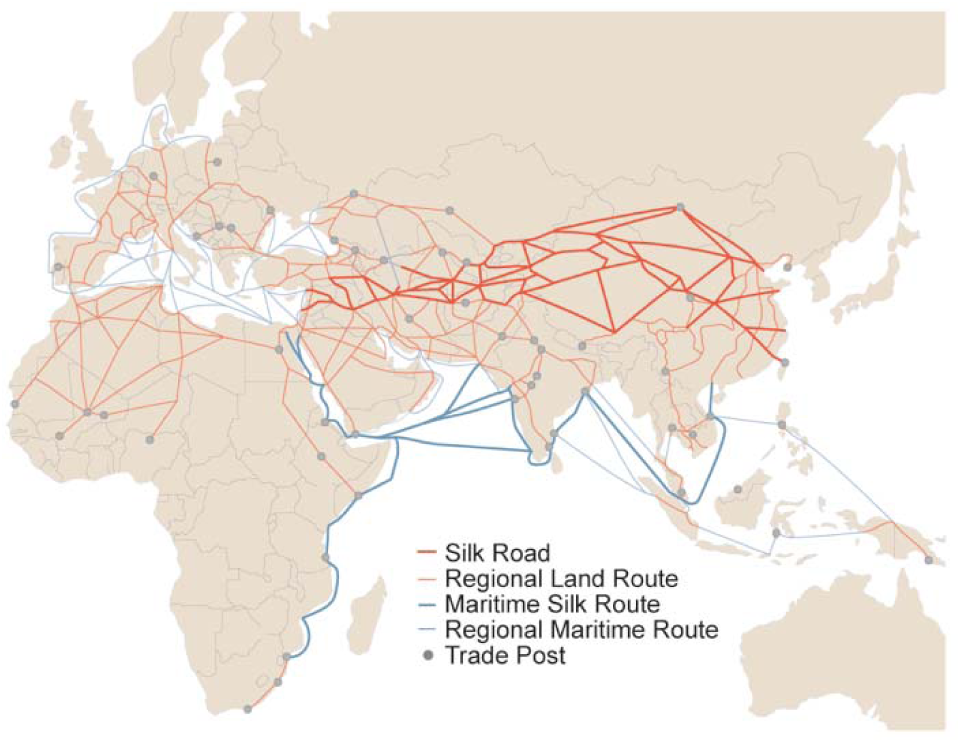
Trade routes active throughout Europe, Africa and Asia by 1400 CE. Nodes (trade cities, oases, and caravanserai) and arcs (the routes between nodes) are from the Old World Trade Routes Project (www.ciolek.com/owtrad.html, accessed February 17, 2016) and are visualized with ArcGIS.

The interval of our timing estimate for the initial migration overlaps with the so-called Middle Asian Interaction sphere in The Age of Integration (2600-1900 BCE), which is marked by increased cultural exchange and trade between civilizations of Egypt, Mesopotamia, the Arabian peninsula, and the Indus Valley (Vogt 1996; Zarins 1996; Parkin and Barnes 2002; Ray 2003; Coningham and Young 2015). East-West contact and trade across the Indian Ocean intensified in the first millennium BCE, when maritime networks expanded to include the eastern Mediterranean, the Red Sea, and the Black Sea (Dilke 1985; Boussac et al. 1995; Ray et al. 1996; Salles 1996). Historical data from the Roman era indicate that crews on trading ships crossing the Indian Ocean comprised fluid assemblages of individuals from diverse regions, brought together under conditions favorable for the transmission of TB (André and Filliozat 1986; Begley and De Puma 1991; Wink 2002; Rauh 2003). These ships would have been an efficient means of spreading *M.tb* among the distant regions involved in trade.

L2 may similarly have an origin in East-West maritime trade across the Indian Ocean, as we infer it arose from a migration event from Eastern Africa to South Eastern Asia during the 1^st^ millennium BCE. In this era, increased sophistication in ship technology allowed for longer voyages (Kent 1979; Blench 1996; Ray et al. 1996; Parkin and Barnes 2002; Wink 2002; Ray 2003). L2 appears to have spread out of Southeast Asia, a highly connected region in our analyses of *M.tb* migration, and is currently found across Eastern Eurasia and throughout South Eastern Asia (fig. 1, fig. 4, fig. S6, fig. S11). Interestingly, although L2 is dominant in Eastern Asia, the region did not appear to have played a prominent role in dispersal of this lineage, except in its exchanges with South Eastern Asia. A recently published study found that the extant *M.tb* population in China traces to a limited number of introductions (Liu et al. 2018), which is consistent with our findings of relatively few exchanges of *M. tb* between Eastern Asia and other regions.

L3 appears to have had relatively low rates of migration throughout its history (fig. 3). The contemporary geographic range of L3 is also narrower, extending east from Northern Africa through Western Asia to the Indian subcontinent (fig. 1). A study of lineage prevalence in Ethiopia showed that L3 is currently concentrated in the north of the country (Comas et al. 2015), consistent with our observed north to south gradient in its distribution on the African continent. This is in opposition to L1, which has a southern predominance in Ethiopia and across Eastern Africa (fig. 1). We estimate L3 emerged in Southern Asia ca. 520 CE (177-739 CE).

Pakistan harbors diverse strains belonging to L3 (fig. S9), and the Southern Asia region was highly connected with Eastern Africa in our analyses (fig. S11). Trade along the Silk Road connecting Europe and Asia was very active in the middle of the first millennium, when we estimate L3 emerged (Hansen 2012; Ball 2016); its distribution suggests it spread primarily along trading routes connecting Northeast Africa, Western Asia, and South Asia (André and Filliozat 1986; Sartre 1991; Hansen 2012; Ball 2016) (fig. 6). We speculate that this occurred *via* overland routes, which may have limited the migration of L3 relative to maritime dispersal of the other lineages.

The geographic distribution of L4 is strikingly broad (fig. 1) and it exhibits minimal population structure (table 1). This suggests L4 dispersed efficiently and continued to mix fluidly among regions, a pattern we would expect if it was carried by an exceptionally mobile population of hosts. L4 is currently concentrated in regions bordering the Mediterranean, and elsewhere throughout Africa and Europe (fig. 1). We estimate the MRCA of L4 emerged in the 1^st^ century CE (range 368 BCE-362 CE), during the peak of Roman Imperial power across the entire Mediterranean world and expansionist Roman policies into Africa, Europe, and Mesopotamia (Luttwak 1976; Isaac 2004). The empire reached its greatest territorial extent in the early second century CE, when all of North Africa, from the Atlantic Ocean to the Red Sea, was under a single power, with trade on land and sea facilitated by networks of stone-paved roads and protected maritime routes (Luttwak 1976; Millar 1993; Ball 2016). Primary sources from Roman civilization attest to trade with China, purposeful expeditions for exploration, cartography, and trade in the Red Sea and Indian Ocean (Pfister and Bellinger 1945; Dilke 1985; Begley and De Puma 1991; Erdkamp 2002; Butcher 2003).

We hypothesize that the broad distribution of L4 reflects rapid diffusion from the Mediterranean region along trade routes extending throughout Africa, the Middle East, and on to India, China, and South Eastern Asia. High rates of migration appear to have been maintained for this lineage over much of its evolutionary history (fig. 3); patterns of connectivity implicate Europe and Africa in its dispersal (fig. S11). The association of L4 with European migrants is well described, particularly migrants to the Americas (Gagneux et al. 2006; Pepperell et al. 2011; Brynildsrud et al. 2018). Here we note bacterial population growth preceded geographic range expansion in L4 ~ca. 15^th^ century (fig. 3), which coincides with the onset of the ‘age of exploration’ (□Ālam and Subrahmanyam 2009) that would have provided numerous opportunities for spread of this lineage from Europeans to other populations. We also note the origin and concentration of this lineage on the African continent. Our sample of L4 isolates includes several deeply rooting African isolates, and African isolates are interspersed throughout the phylogeny (fig. 4, fig. S6, fig. S8).

The migratory histories of L5, L6, and L7 are less complicated than those of lineages 1-4. Specifically, L5 and L6 are restricted to Western Africa and L7 is found only in Ethiopia (fig. 4, fig. S6). The reasons for the restricted distributions of these lineages are not immediately obvious: there is evidence in our analyses that other lineages migrated in and out of Western Africa, and Eastern Africa emerged as highly connected and central to the dispersal of *M.tb* (fig. 5). A potential explanation is restriction of the pathogen population to human sub-populations with distinct patterns of mobility and connectivity that did not facilitate dispersal. This is likely the case for L7, which was discovered only recently (Blouin et al. 2012), and is currently largely restricted to the highlands of northern Ethiopia (Firdessa et al. 2013; Comas et al. 2015). In the case of L6 (also known as *Mycobacterium africanum*), there is evidence suggesting infection is less likely to progress to active disease than for *M. tuberculosis sensu stricto* (Jong et al. 2008), which could have played a role in limiting its dispersal.

Our reconstructions of *M.tb’*s migratory history suggest that patterns of migration were highly dynamic: the pathogen appears to have dispersed efficiently, in complex patterns that nonetheless preserved the distinct structure of each lineage. Some findings, notably inference of population expansion, were consistent across lineages. Though growth of the global *M.tb* population has been described previously (Comas et al. 2013; Pepperell et al. 2013), our results here suggest that the pace and magnitude of expansion, and its apparent relationship to trends in migration, varied among lineages (fig. 3, fig. S3, fig. S11).

Our analyses suggest that the expansion of L2 was preceded by an impressive increase in its rate of migration (fig. 3), implying that growth of the pathogen population was facilitated by expansion into new niches. Our phylogeographic reconstructions implicate Russia, Central Asia, and Western Asia in the recent migratory history of L2 (fig. S10, fig. S11), which is consistent with a published phylogeographic analysis of L2 (Luo et al. 2015). The inferred timing of the growth and increased migration of L2 (~ca. 13^th^ century) is close to the well documented incursion of *Yersinia pestis* from Central Asia into Europe that resulted in explosive plague epidemics (Benedictow 2004). The experience with plague suggests that patterns of connectivity among humans and other disease vectors were shifting at this place and time, which would potentially open new niches for pathogens including *M.tb*.

We estimate that L1 underwent expansion ~ca. 17^th^ century (fig. 3) but in this case it appears to have grown *in situ*, e.g. due to changing environmental conditions such as increased crowding, and/or growth of local human populations. A study of the molecular epidemiology of TB in Vietnam identified numerous recent migrations of L2 and L4 into the region, versus a stable presence of L1 (Holt et al. 2018); this is consistent with our finding of higher recent rates of migration for L2 and L4 versus L1 (fig. 3). A pattern similar to L1 has been identified previously, in the delay between dispersal of *M.tb* from European migrants to Canadian First Nations and later epidemics of TB driven by shifting disease ecology (Pepperell et al. 2011).

These results demonstrate the complex relationship between *M.tb* population growth and migration, and show that under favorable conditions the pathogen can expand into novel niches or accommodate growth in an existing niche.

In a previous study, analyses of synonymous and non-synonymous SFS have been used to delineate effects of purifying selection, linkage of sites, and population expansion on global populations of *M.tb* (Pepperell et al. 2013). Simulation studies have shown that purifying selection can affect demographic inference with BEAST and SFS-based methods (Ewing and Jensen 2015; Lapierre et al. 2016). Although our analyses here using ∂a∂i were restricted to synonymous SFS, it is likely that inference of population size changes with this method and with BEAST were affected by purifying selection on this fully linked genome. The magnitude of inferred expansions may thus reflect both population size changes and background selection, and should not be interpreted as direct reflections of historical changes in census population size. We did not detect an effect of purifying selection on inference of migration in our three population simulation analyses (note S2, fig. S12, fig. S13), but differences in the strength of purifying selection could contribute to the lineage-specific differences we observed in the size of inferred population expansions: i.e., genome-wide patterns of purifying selection could differ among lineages. Previous evidence has suggested that the fitness trade-offs of drug resistance mutations vary among lineages (Mortimer et al. 2018), making this intriguing possibility potentially feasible.

This study has some important limitations. We did not attempt to estimate the rate or timescale of *M.tb* evolution, instead relying on published rates that were calibrated with ancient DNA. This is an active area of research, and newly discovered ancient *M.tb* DNA samples will likely refine inference of both the timing and locations of historical migration events, though it is critical to note that recent substitution rate estimates of *M.tb* have converged on rates around 5×10^−8^ substitutions per site per year (Eldholm et al. 2016). Even when substitution rate estimates can be estimated with confidence, the precision with which individual events can be dated using genetic data should not be over-stated, as evidenced by broad 95% credible intervals for internal node date estimates (e.g., Eldholm et al. 2016). Our goal here was to reconstruct historical migration of *M.tb* throughout Eurasia and Africa and place this evolutionary history within a broad historical context; the historical phenomena that we connect with the spread of TB involved vast areas and extended over hundreds and in some cases thousands of years. Our reconstruction of the global dispersal of TB within a temporal framework provided by ancient *M.tb* DNA analysis links spread of the disease to the first ~1500y of the common era, a period of remarkable intensification in the connectedness among peoples of Africa, Asia and Europe (Green 2018).

## Methods

### Lineage Frequencies

The SITVIT WEB database (Demay et al. 2012), which is an open access *M.tb* molecular markers database, was accessed on September 5, 2016. Spoligotypes were translated to lineages based on the following study (Shabbeer et al. 2012). The following conversions were also included: EAI7-BGD2 for L1, CAS for L3, and LAM7-TUR, LAM12-Madrid1, T5, T3-OSA, and H4 for L4. Isolates containing ambiguous spoligotypes (denoted with >1 spoligotype) were inspected manually and assigned to appropriate lineages. Relative lineage frequencies of lineages 1-6 for each country containing data for >10 isolates were calculated and plotted with the rworldmap package in R (South 2016).

### Sample Description

#### Old World collection

We assembled/aligned publicly available whole genome sequences (WGS) of thousands of *M.tb* isolates from recently published studies and databases for which country of origin information were known and fell within regions traditionally defined as the Old World. Isolates were assembled via reference guided assembly (RGA) when FASTQ data were available and by multiple genome alignment (MGA) when only draft genome assemblies were accessible (see below). As we were interested in reconstructing historical migrations of the pathogen, we excluded countries where the majority of contemporary TB cases are identified in recent immigrants (Government of Canada 2005; White et al. 2017; Australian Government Department of Health and Ageing; Centers for Disease Control; Institute of Environmental Science and Research Limited; Public Health England). Due to computational limitations (BEAST analyses), we necessarily took measures to limit our dataset to <600 isolates. For countries with large numbers of available genomes, we implemented a sub-sampling strategy similar one previously described (Thorpe et al. 2017), whereby phylogenetic lineage diversity was captured thus minimizing the overrepresentation of clonal complexes (e.g., outbreaks): phylogenetic inference on all isolates available from a country was performed with Fasttree (Price et al. 2010) and a random isolate was selected from each clade extending from *n* branches, where *n* was the desired number of isolates from the country. Numbers of isolates per country were selected based on the availability of appropriate genome sequence data as well as relative TB prevalence (fig. S1) (World Health Organization 2017). All isolates belonging to lineages 5-7 were retained. As a whole, this dataset reflects a ‘mixed’ sampling scheme (Lapierre et al. 2016), where lineages L5-L7 are overrepresented relative to their contemporary frequencies (fig. 1). At the lineage-specific scale, L1-L4 approximate random sampling of available genomes. Our final Old World collection consisted of the WGS of 552 previously published *M.tb* isolates collected from 51 countries spanning 13 UN geoscheme subregions. Accession numbers and pertinent information about each sample can be found in table S1.

We note that our sample necessarily contains a large number of drug-resistant isolates as these are more commonly sequenced. We also acknowledge that the studies we draw genomes from may have been subject to other sampling biases for which we are unaware.

#### Northern and Central American collection

For one analysis, we included an additional 15 isolates from a previous study (Comas et al. 2015) for which the country of origin was within the Americas. Isolates were assembled via RGA (see below) and their genotypes at the 3,838,249 bp considered for all analyses of the Old World collection were extracted.

### Reference Guided Assembly

Previously published FASTQ data were retrieved from the National Center for Biotechnology Information (NCBI) sequence read archive (SRA) (Leinonen et al. 2011). Low-quality bases were trimmed using a threshold quality of 15, and reads resulting in less than 20bp length were discarded using Trim Galore! (http://www.bioinformatics.babraham.ac.uk/projects/trim_galore/), which is a wrapper tool around Cutadapt (Martin 2011) and FastQC (http://www.bioinformatics.babraham.ac.uk/projects/fastqc/). Reads were mapped to H37Rv (NC_000962.3) (Cole et al. 1998) with the MEM algorithm (Li 2013). Duplicates were removed using Picard Tools (http://picard.sourceforge.net), and local realignment was performed with GATK (DePristo et al. 2011). To ensure only high quality sequencing data were included, individual sequencing runs for which <80% of the H37Rv genome was covered by at least 20X coverage were discarded, as were runs for which <70% of the reads mapped as determined by Qualimap (García-Alcalde et al. 2012). Pilon (Walker et al. 2014) was used to call variants with the following parameters: --variant --mindepth 10 --minmq 40 --minqual 20.

### Multiple Genome Alignment

Draft genome assemblies were aligned to H37Rv (NC_000962.3) (Cole et al. 1998) with Mugsy v1.2.3 (Angiuoli and Salzberg 2011). Regions not present in H37Rv were removed and merged with the reference-guided assembly.

### SNP alignment

Variant calls (VCFs) were converted to FASTAs with in-house scripts that treat ambiguous calls and deletions as missing data (available at https://github.com). Transposable elements, phage elements, and repetitive families of genes (PE, PPE, and PE-PGRS gene families) that are poorly resolved with short read sequencing were masked to missing data. Isolates with >20% missing sites were excluded from the Old World collection (table S1). Variant positions with respect to H37Rv were extracted with SNP-sites (Page et al. 2016) resulting in 60,818 variant sites. Only sites where at least half of the isolates had confident data (i.e., non-missing) were included in the phylogeographic models and population genetic analyses (60,787 variant sites; 3,838,249 bp). 1.7% of variant sites landed in loci associated with drug resistance (table S3).

### Geographic Information

Geographic locations for each of the 552 samples in the Old World collection were obtained from NCBI and/or the publications in which the isolates were first described. When precise geographic information was available (e.g., city, province, etc.), coordinates were obtained from www.mapcoordinates.net. When only country level geographic information was available, the ‘Create Random Point’ tool in ArcGIS 10.3 was used to randomly place each isolate without specific latitude and longitude inside its respective country; inhospitable areas (e.g., deserts and high mountains) and unpopulated areas from each country using 50m data from Natural Earth (http://www.naturalearthdata.com/downloads, accessed February 17, 2016) were excluded as possible coordinates. The ‘precision’ column of table S1 reflects which method was used.

### Trade Route Information

Data for all trade routes active throughout Europe, Africa, and Asia by 1400 CE were compiled from the Old World Trade Routes (OWTRAD) Project (www.ciolek.com/owtrad.html, accessed February 17, 2016). For each route, both node information (trade cities, oases, and caravanserai) and arc information (the routes between nodes) were imported into ArcGIS (fig. 6). *M.tb* isolate locations were also imported as points and the ‘Generate Near Table’ tool was used to assign each isolate to its nearest node in the trade network and is listed in the ‘NearPost’ column of table S1.

### Maximum Likelihood Inference

We used RAxML v8.2.3 (Stamatakis 2014) for maximum likelihood phylogenetic analysis of the Old World collection (all sites where at least half of isolates had non-missing data) under the general time reversible model of nucleotide substitution with a gamma distribution to account for site-specific rate heterogeneity. Rapid bootstrapping of the corresponding SNP alignment was performed with the -autoMR flag, converging after 50 replicates. Tree visualization was created with the ggtree package in R (Yu et al. 2017).

### Phylogeographic & Demographic Inference with BEAST

The Old World collection SNP alignment and individual lineage SNP alignments were analyzed using the Bayesian Markov Chain Monte Carlo coalescent method implemented in BEAST v1.8 (Drummond and Rambaut 2007) with the BEAGLE library (Ayres et al. 2012) to facilitate rapid likelihood calculations. Analyses were performed using the general time reversible model of nucleotide substitution with a gamma distribution to account for rate heterogeneity between sites, a strict molecular clock, and both constant and Bayesian skyline plot (BSP) demographic models. Country of origin or the UN subregion for each isolate was modeled as a discrete phylogenetic trait (Lemey et al. 2009). All Markov chains were run for at least 100 million generations, sampled every 10,000 generations, and with the first 10,000,000 generations discarded as burn-in; replicate runs were performed for analyses and combined to assess convergence. Estimated sample size (ESS) values of non-nuisance parameters were >200 for all analyses. Site and substitution model choice were based on previous analyses of *M.tb* global alignments as opposed to an exhaustive comparison of models which would require unreasonable computational resources. Strict vs relaxed molecular clocks did not result in altered trends of migration at the lineage level, and comparisons between analyses using strict and relaxed clocks show strong correlation between the estimated height of nodes (e.g., R^2^ > 0.97; fig. S14). Table S4 provides a summary of BEAST analyses presented and the results derived from them. Tree visualizations were created with FigTree (http://tree.bio.edu.ac.uk/software/figtree/) and the ggtree package in R (Yu et al. 2017).

### Phylogeographic reconstruction: limitations and alternatives

These phylogeographic reconstructions are clearly sensitive to sampling, since we cannot identify the roles of unsampled regions in *M.tb’*s migratory history. We maximized geographic diversity in our sample, but were limited by available data and some regions – notably Middle Africa, Northern Africa, and Western Asia – are absent or underrepresented in our sample (fig. S1). Defining the contributions of these undersampled regions to *M.tb’*s migratory history awaits more samples and/or further method development.

De Maio *et al*. (2015) note the sensitivity of discrete trait phylogeographic inference in BEAST to sample selection, as well as overconfidence in the precision of geographic inference, and propose BASTA as an alternative (De Maio et al. 2015). BASTA is sensitive to the choice of prior and we did not have ancillary data to guide the selection of a prior for the Old World migratory history of *M.tb*, precluding its use here. We investigated ∂a∂i as an alternative tool for phylogeographic inference but it did not perform well for this application under conditions of complete linkage of sites (note S3, fig. S15, fig. S16, table S5, table S6). The phylogeographic inference method implemented here relies on the assumption that sample size reflects deme size (Lemey et al. 2009; De Maio et al. 2015), and within the constraints of available data, we attempted to adjust our sample sizes according the regional prevalence of TB (see *Methods* and fig. S1). We also interrogated the relationship between regional sample size and inferred migration rate and did not observe a strong correlation (fig. S17). According to the classifications proposed by Lapierre *et al*. (2015), our Old World collection represents a ‘mixed’ sampling scheme (see *Methods*).

### Demographic inference from the observed site frequency spectrum (SFS)

SNP-sites (Page et al. 2016) was used to convert the Old World collection alignment to a multi-sample VCF and SnpEff (Cingolani et al. 2012) was used to annotate variants with respect to H37Rv (NC_000962.3) (Cole et al. 1998) as synonymous, non-synonymous, or intergenic. Loci at which any sequence in the population had a gap or unknown character were removed from the data set. Demographic inference with the synonymous SFS for each of the seven lineages and the entire collection was performed using ∂a∂i (Gutenkunst et al. 2009). We modeled constant population size (standard neutral model), an instantaneous expansion model, and an exponential growth model, and identified the best-fit model and maximal likelihood parameters of the demographic model given our observed data. Our parameter estimates, ν and τ, were optimized for the instantaneous expansion and exponential growth models. Uncertainty analysis of these parameters were analyzed using the Godambe Information Matrix (Coffman et al. 2016) on 100 samplings of the observed synonymous SFS with replacement and subsequent model inference.

### Population genetic statistics

Nucleotide diversity (π) and Watterson’s theta (□) for various population assignments (e.g., lineage, UN subregion) were calculated with EggLib v2.1.10 (De Mita and Siol 2012).

### Analysis of Molecular Variance (AMOVA)

AMOVAs were performed using the ‘poppr.amova’ function (a wrapper for the ade4 package (Dray et al. 2007) implementation) in the poppr package in R (Kamvar et al. 2014). Bins were assigned via the following classification systems: UN geoscheme subregions and Level 1 (‘botanical continents’) of the World geographical scheme for recording plant distributions. Isolate assignation can be found in table S1. Genetic distances between isolates were calculated with the ‘dist.dna’ function of the ape v4.0 package in R (Paradis et al. 2004) from the SNP alignment of the Old World collection.

### Mantel tests

Great circle distances between *M.tb* isolate locations were calculated with the ‘distVincentyEllipsoid’ function in the geosphere R package (Hijmans et al. 2016). Geographic distances between isolate locations along the trade network were calculated by adding the great circle distances from the isolates to the nearest trade hubs and the shortest distance between trade hubs along the trade network; the latter was determined using an Origin-Destination Cost Matrix and the ‘Solve’ tool in the Network Analyst Toolbox of ArcGIS which calculates the shortest distance from each origin to every destination along the arcs in the trade network. In the event that two isolates were assigned to the same trade post, the great circle distance between the isolates was used. To calculate the geographic distance between isolates in a manner that reflects human migrations, the great circle distance between isolates and waypoints were summed. These were calculated with a custom R function (available at https://github.com) using a series of rules to define whether or not the path between isolates would have gone through a waypoint. For all three distance metrics, values were log transformed and standardized. Genetic distances between isolates were calculated with the ‘dist.dna’ function in the ape v4.0 package in R (Paradis et al. 2004) from the SNP alignment. The ‘mantel’ function of the vegan package in R (Oksanen et al. 2017) was used to perform a Mantel test between the genetic distance matrix and each of the three geographic matrices for both the Old World collection and each individual lineage. Four of the 552 isolates were excluded from these analyses as they were from Kiribati and trade networks spanning this region were not compiled.

### Relationship between genetic diversity and geographic distance from Addis Ababa

For this analysis, we added Northern and Central American datasets, assembled in an identical manner to those of the Old World collection and masked at sites where less than half of the Old World collection had confident data (3,838,249 bp). For each UN subregion, the mean latitude and longitude coordinates for all *M.tb* isolates within the region were calculated. The great circle distances from these average estimates for regions to Addis Ababa were then calculated, using waypoints for between-continent distance estimates to make them more reflective of presumed human migration patterns (Ramachandran et al. 2005). Cairo was used as a waypoint for Eastern Europe, Central Asia, Western Asia, Southern Asia, Eastern Asia, and South Eastern Asia; Cairo and Istanbul were used as waypoints for Western Europe and Southern Europe; Cairo, Anadyr, and Prince Rupert were used as waypoints for Northern and Central America. The distance between each region and Addis Ababa were the sum of the great circle distances between the two points (the average coordinates for the UN subregion and Addis Ababa) and the waypoint(s) in the path connecting them, plus the great circle distance(s) between waypoints if two were used. Treating each UN subregion as a population, the relationship between genetic diversity (assessed with π) and geographic distance from Addis Ababa were explored with linear regression for both the entire Old World collection and individual lineages in R (R Development Core Team). Code is available at https://github.com.

### Migration Rate Inference

Migration rates through time were inferred from the Bayesian maximum clade credibility trees for the entire Old World collection of *M.tb* isolates (*n =* 552). Individual lineages that contain isolates from multiple UN subregions (i.e., L1: *n =* 89, L2: *n =* 181, L3: *n = 65*, and L4: *n =* 143) were extracted and plotted separately. Only nodes with posterior probabilities greater than or equal to 80% were considered. A migration event was classified as a change in the most probable reconstructed ancestral geographic region from a parent to child node. Median heights of the parent and child nodes were treated as a range of time that the migration event could have occurred. The rate of migration through time for each lineage or the Old World collection was inferred by summing the number of migration events occurring across every year of the time-scaled phylogeny, divided by the total number of branches in existence during each year of the time-scaled phylogeny (both those displaying a migration event and those that do not). Code for these analyses is available at https://github.com.

Additionally, relative migration rates between UN subregions were derived from the BEAST analyses of phylogeography. The Bayesian stochastic search variable selection method (BSSVS) for identifying the most parsimonious migration matrix implemented in BEAST as part of the discrete phylogeographic migration model (Lemey et al. 2009) allowed us to use Bayes factors (BF) to identify the migration rates with the greatest posterior support and provide posterior estimates for their relative rates. Strongly supported relative rates (BF > 5) and connectivity among subregions were visualized with Cytoscape v3.2.0 (Shannon et al. 2003) and superimposed onto a map generated with the ‘rworldmap’ package in R (South 2016).

### Effect of selection on estimates of migration

We performed demographic forward-in-time simulations using the SFS_CODE package (Hernandez 2008), which allows for demographic models with arbitrarily complex migration and selection regimes. Our simulations were performed under a simple two population model or with a more complex three population model. In all simulations, *N*_*e*_ for each population was 1000, □ was 0.001 (O’Neill et al. 2015), and migration between each pair of populations was symmetrical. As there is substantial evidence for little to no recombination in the *M.tb* genome, our simulations were performed without recombination.

The two population simulations were performed under three scenarios: 1) no migration between populations after initial divergence; 2) constant migration after divergence (per generation *M* = 0.5) without selection; and 3) constant migration (*M* = 0.5) with purifying selection (25% of alleles of each population have a population selection coefficient of −1.0, and the rest are neutral) after divergence.

The three population simulations were performed under five scenarios: 1) no migration between populations after simultaneous divergence of the three populations; 2) constant, symmetrical migration after divergence (per generation *M* = 0.5 for all population pairs) without selection; 3) constant, symmetrical migration (*M* = 0.5) with purifying selection (25% of alleles in all populations have a population selection coefficient of −1.0, and the rest are neutral); 4) constant, asymmetrical migration after divergence (*M* = 0.5 for migration between pop0 and pop1, *M* = 5.0 for migration between pop1 and pop2, and *M* = 0 for migration between pop0 and pop2) without selection; and 5) constant, asymmetrical migration after divergence (*M* = 0.5 between pop0 and pop1, *M* = 5.0 between pop1 and pop2, and *M* = 0 between pop0 and pop2) with purifying selection (25% of alleles in all populations have a population selection coefficient of −1.0, and the rest are neutral).

For all simulations, 25 samples were taken from each population, and sequences of 100000 bases were generated. Twenty simulations were performed under each scenario for both the 2 population (60 simulations) and 3 population (100 simulations) models. Each sequence alignment was subsequently subjected to migration analysis in ∂a∂i (Gutenkunst et al. 2009, see note S2) and BEAST v1.8.4 (Drummond and Rambaut 2007). For each Bayesian coalescent analysis, the HKY+G substitution model, a constant population model, and a strict molecular clock model were used. A discrete symmetrical migration model (Lemey et al. 2009) was used to determine migration rates, and BSSVS (Lemey et al. 2009) was used to estimate BF support for migration rates in the 3 population simulations. All Markov chains were run for 10 million generations or until convergence, with samples taken every 10,000 steps, and 10% discarded as burn-in. The package SpreaD3 v0.96 (Bielejec et al. 2016) was used to calculate BF support for migration rates.

