## supplemental information for "Lineage specific histories of *Mycobacterium tuberculosis* dispersal in Africa and Eurasia"

### Note S1. Supplementary note about the relationship between genetic diversity and geographic distance from Addis Ababa.

Our finding of no significant decline in *M.tb* diversity as a function of distance from Addis Ababa conflicts with a previously published report (Comas et al. 2015). To ensure our differing results were not driven by a lack of samples from the Americas, we repeated the analysis including the same samples from the Americas used in Comas *et al*. (2015), and obtained results that were similar to the Old World collection alone (adjusted R-squared = 8.9 x 10^-4^, *p* = 0.34 versus adjusted R-squared = -0.1, *p* = 0.88, respectively). We did not find a trend in diversity as a function of distance at the lineage specific level either (fig. S5, table S2), and speculate that the significance obtained by Comas *et al*. (2015) may have been driven by the lineage makeup of their defined regions (i.e., samples from the Americas consisted solely of L4 isolates while other regions harbored isolates from multiple lineages) and/or the larger number of isolates from Ethiopia relative to other regions. Our analysis using a larger sample size, finer geographic resolution, and the use of waypoints of human migration in calculating distances from Addis Ababa do not lend support to the hypothesis of serial bottlenecks related to out of Africa migrations having shaped diversity of non-African populations of *M.tb*.

### Note S2. Supplementary note about the effects of selection on inference of migration.

The effects of population expansion, linkage, and purifying selection on *M.tb* genetic diversity have previously been demonstrated (Pepperell et al. 2013). Given these previous observations, we were curious about a potential impact of purifying selection on inference of migration. To address this question, we simulated data under demographic models with and without selection and migration, and then analyzed the resulting sequence alignments in BEAST. Results of our three population model suggested that purifying selection had a statistically negligible effect on migration rates, which can be observed from plots of the mean relative rates (fig. S12) or of the relative support of migration rates (fig. S13). We note that the discrete migration model implemented in BEAST was able to capture much of the asymmetry of our three population asymmetrical simulations as evidenced by the distribution of relative migration rates and Bayes factor (BF) support for said rates. BEAST also consistently produced similar BF support for rates estimated from data simulated under symmetrical migration models (i.e., those with global *M* = 0.5 or 0.0). Our simulations thus suggest that consistent purifying selection is unlikely to dramatically affect estimates of, or support for, migration rates between populations in these scenarios.

### Note S3. Supplementary note about migration inference with ∂a∂i.

We attempted to use ∂a∂i to infer *M.tb* migration history as a complement to our analyses performed in BEAST. Briefly, EasySFS was used to convert the multi-sample VCF of L1 to a two-dimensional SFS (<https://github.com/isaacovercast/easySFS>). Populations were defined as India and the rest of the world (RoW) and projected from *n*=31,58 to *n*=25,25 (India and RoW, respectively). Migration inference with the synonymous SFS was performed using ∂a∂i. We modeled no split (standard neutral model), a split with no migration, a split with symmetric migration, a split with unidirectional migration (India to RoW), and a split with asymmetric migration and identified the best-fit model and maximal likelihood parameters of the migration model given our observed data. Parameters v1 and v2 were fixed according to their values estimated from a model of instantaneous expansion. Our parameter estimates m and τ, were optimized for each migration model. We used the Akaike information criterion (AIC) estimator for model selection and calculated the Poisson residuals between model and data for each best-fit model.

Results of these analyses indicated that all tested models fit poorly, as indicated by clustering of residuals from the 2D site frequency spectra, and the method was therefore unreliable for model selection (fig. S15, fig. S16, table S3).

We also used ∂a∂i to analyze simulated data for a fully linked genome under simple models of migration (see *Methods* for details of simulations). Briefly, EasySFS was used to convert the multi-sample VCF of simulated populations to a two-dimensional SFS (*n*=25,25). We modeled a split with no migration, a split with symmetric migration, a split with unidirectional migration (pop0 to pop1), and a split with asymmetric migration. We identified the best-fit model and maximal likelihood parameters of the migration models across replicates for each simulated condition. Our parameter estimates v1, v2, m and τ, were optimized for each migration model. We used the AIC estimator for model selection.

Results of these analyses also indicated that the methods performed poorly in this context, with the correct model only inferred in 26% of instances (table S4). These results are consistent with prior research indicating that SFS-based methods perform well for fully linked genomes when inference is done under very simple models but not more complex models (Pepperell et al. 2013).

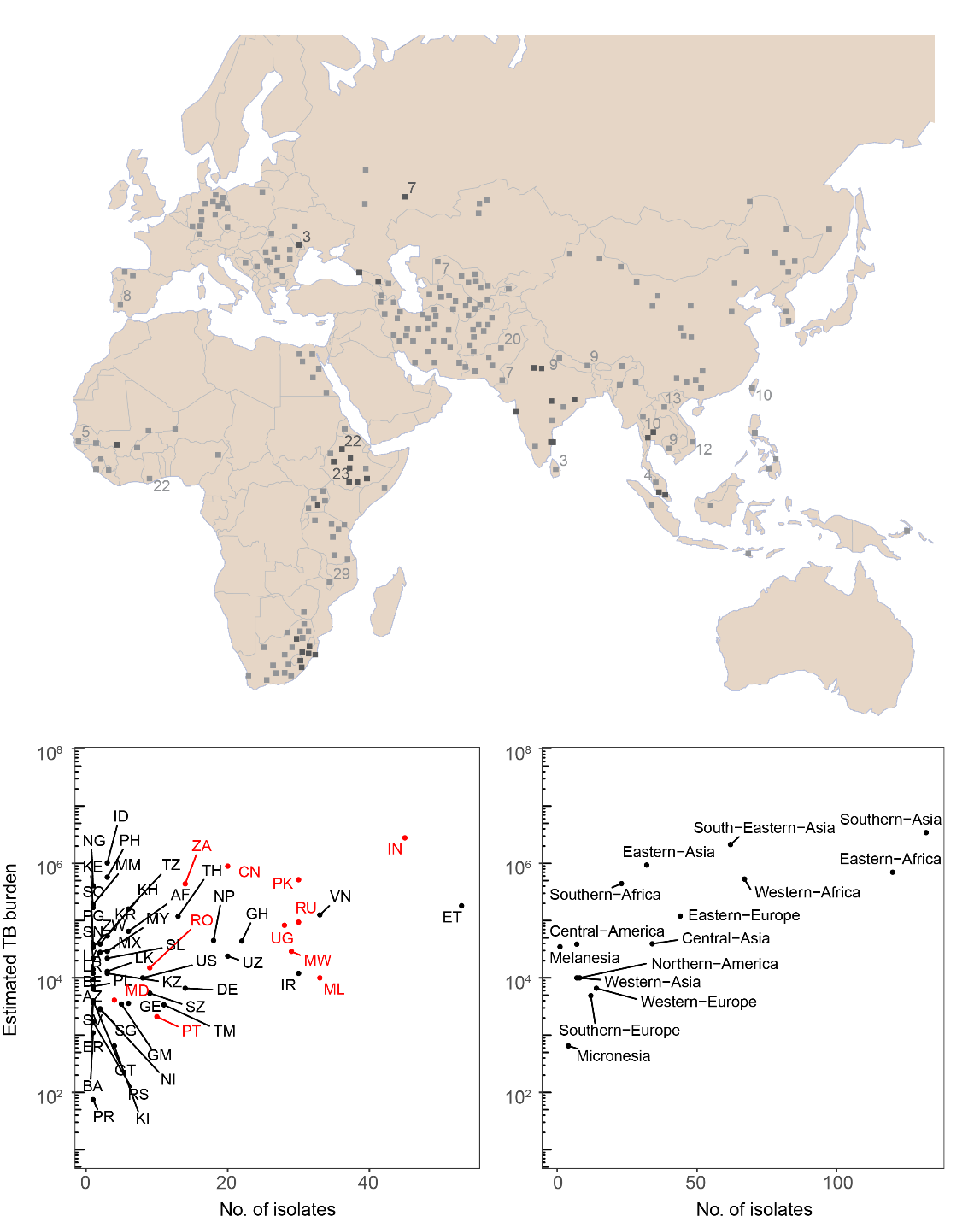

A

B

C

Fig. S1. Sampling strategy. (A) Map of the geographic locations for each of the 552 samples in the Old World collection. Black squares reflect the coordinates of the origin of an isolate when precise geographic information was available (e.g., city), whereas light grey squares reflect the randomized coordinates assigned to an isolate within the hospitable areas of the county from which it originated (see methods). Numbers next to squares designate the number of samples originating from a given location. (B) The number of isolates from each country in the Old World collection versus the estimated TB burden of that country. Estimated TB burden reflects the 2016 TB burden estimates by WHO. Iso2 codes for each country are labelled, and countries for which the total number of available genomes were downsampled (see *Methods*) are colored in red. (C) The number of isolates from each UN sub-region in the Old World collection versus the estimated TB burden for that sub-region. Sub-region TB burdens were determined by summing the estimated TB burden reported for all countries within the sub-region for which we had isolates in our Old World collection.

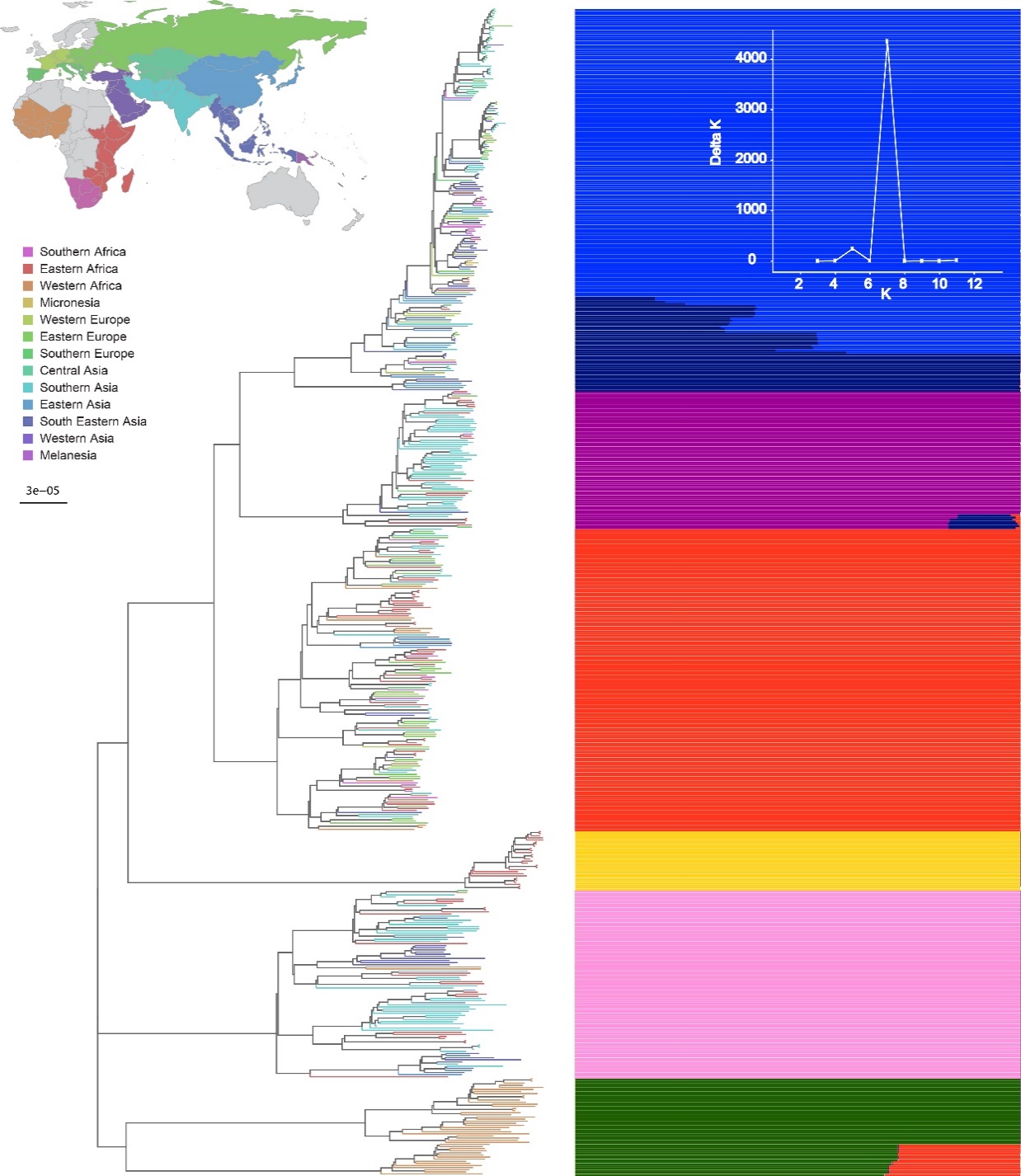

Fig. S2. Maximum likelihood phylogeny of the Old World collection of 552 Mycobacterium tuberculosis isolates. Phylogenetic analysis was performed with RAxML using the general time reversible model of nucleotide substitution under the Gamma model of rate heterogeneity on all sits where at least half of the isolates had confident genotypes (3,838,249bp). Rapid bootstrapping was performed with the -autoMR flag on the corresponding variant sites of the alignment (60,787), converging after 50 replicates. Tip labels are colored with respect to their geographic origin according to the UN geoscheme, as pictured in the legend.

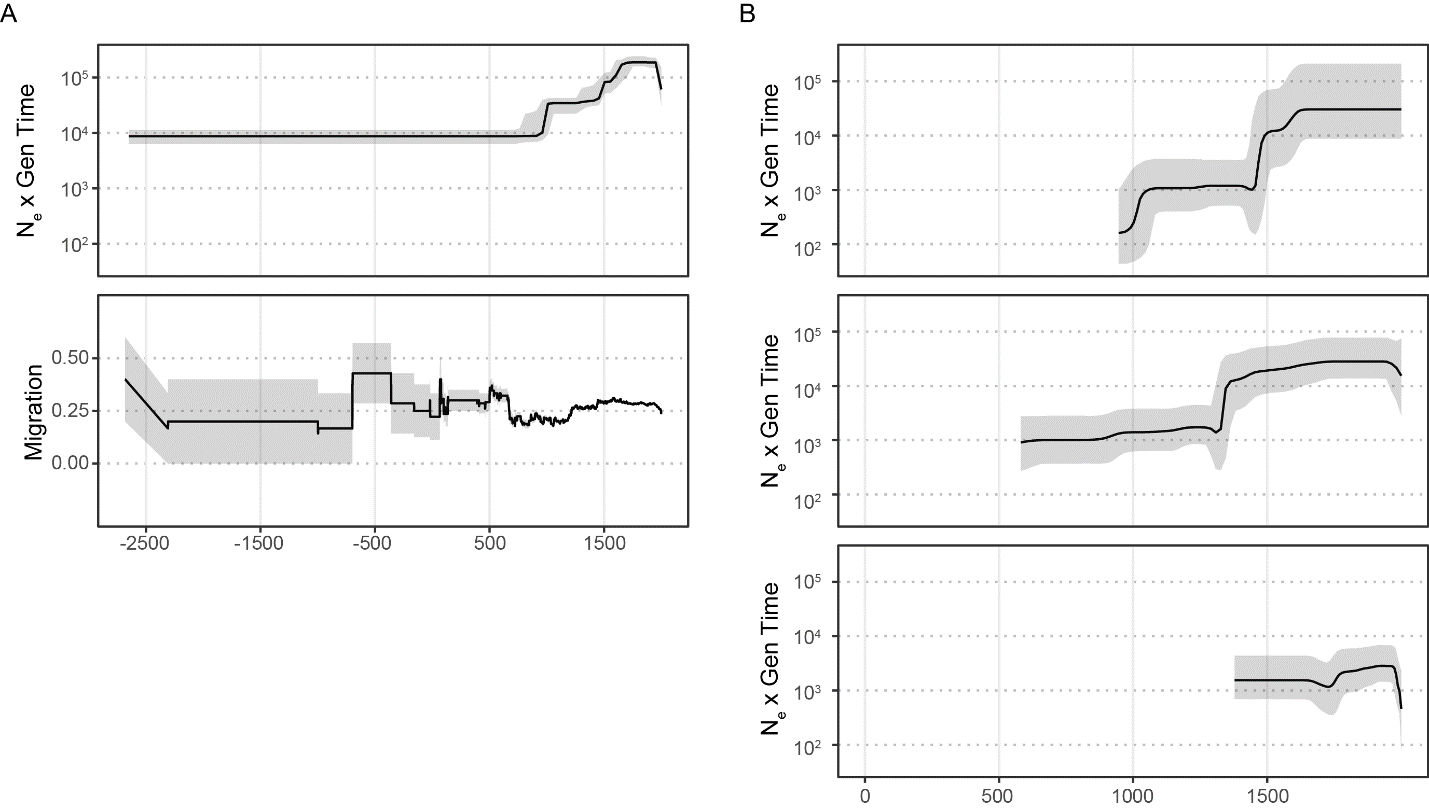

B

A

Fig. S3. Demographic histories and migration rates of *M.tb*. (A) Old World collection. Top panel – Bayesian skyline plot (BSP) shows the inferred change in effective population through time. Black lines denote median N_e_ and gray shading the 95% highest posterior density. Bottom panel - migration rate through time inferred from the phylogeographic analysis (see *Methods*). Grey shading depicts the rates inferred after the addition or subtraction of a single migration event, demonstrating the uncertainty of rate estimates from the early history of the phylogeny. (*B*) BSPs for Lineages 5-7 (top to bottom, respectively). Lineages 5-7 are only found in only one subregion each throughout their phylogenies resulting in a migration rate of zero through time. Dates are shown in calendar years and are based on scaling the phylogeny with a substitution rate of 5 x 10^-8^ substitutions/site/year.

A

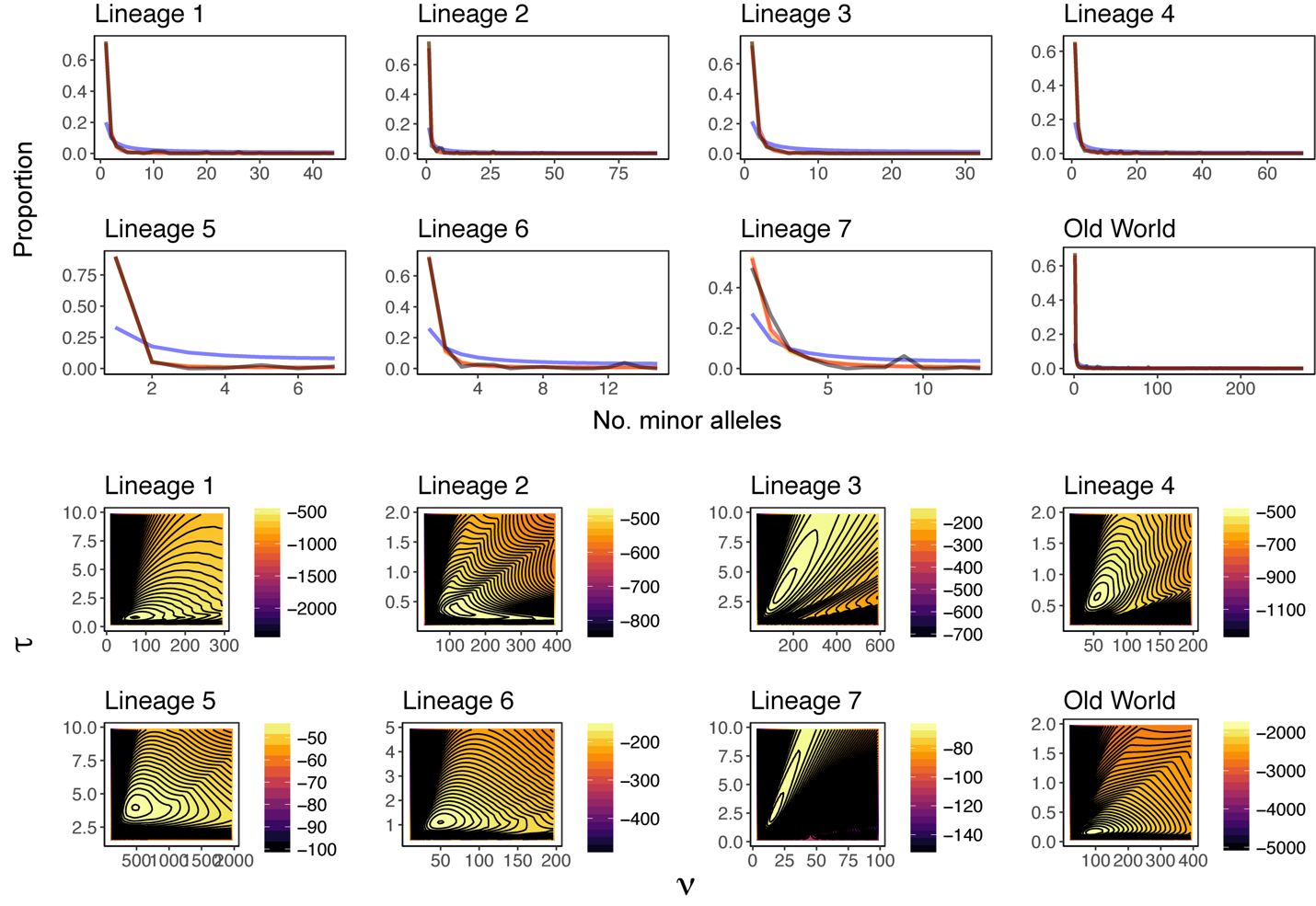

B

Fig. S4. Demographic analyses with ∂a∂i. (A) Folded site frequency spectrums. Demographic inference with the synonymous SFS for the entire Old World collection and each of the seven lineages was performed using methods implemented in ∂a∂i. The observed folded SFS is plotted in black, while the expected folded SFS under constant population size is plotted in blue, instantaneous expansion in red, and exponential growth in orange. (B) Likelihood surfaces. Heatmaps of log10 likelihood values (see scale bars) over a range of values for two demographic parameters in a model of instantaneous expansion are plotted: generations since expansion (τ) and N_e_/N_anc_(ν).

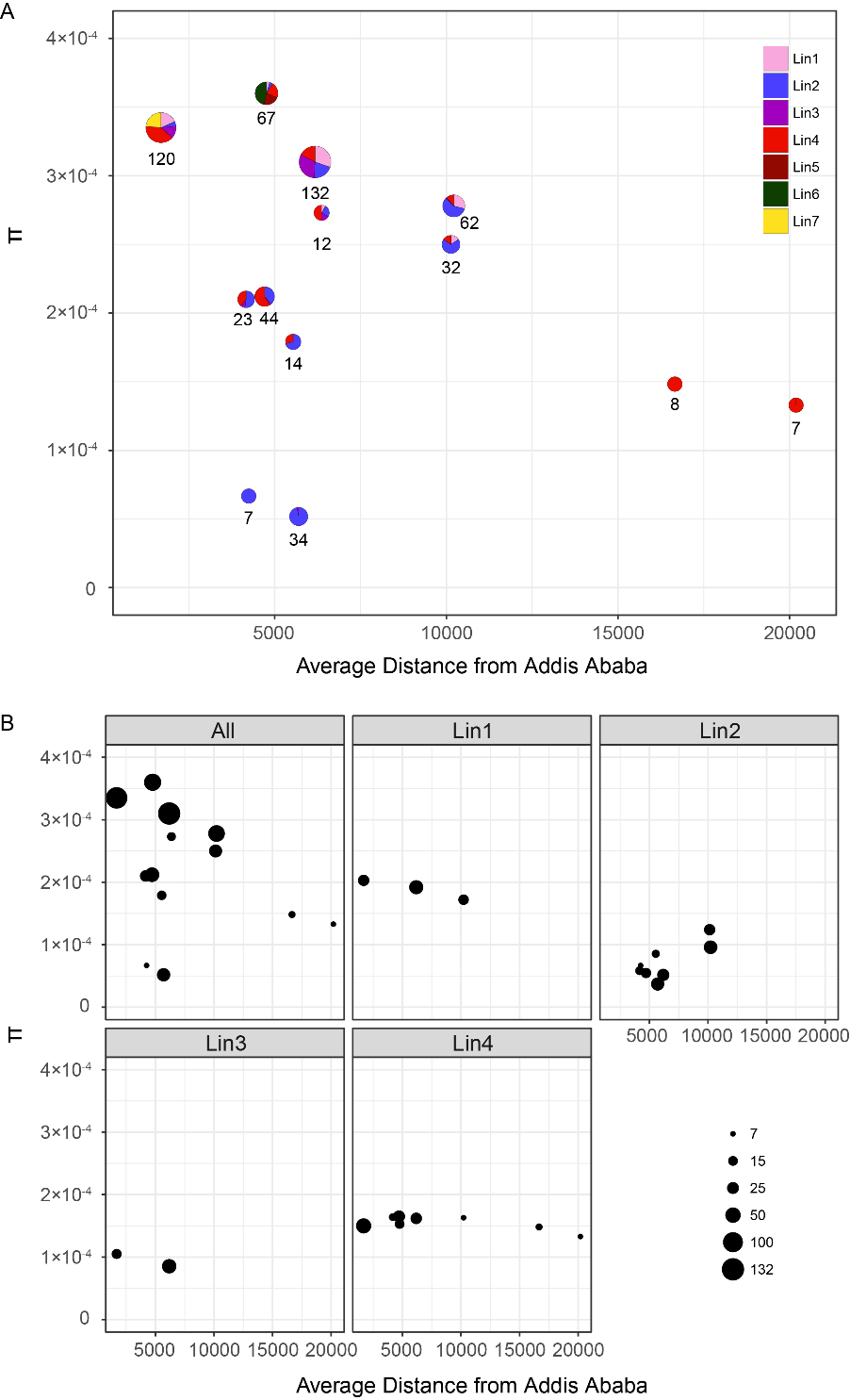

A

B

Fig. S5. Relationship between genetic diversity and geographic distance from Addis Ababa. (A) Diversity as a function of distance for all isolates among 13 UN subregions. Treating each UN subregion as a population, nucleotide diversity (π) was compared to the mean distance of isolates from the region to Addis Ababa (see *Methods*). Point size reflects the number of isolates per subregion and colors reflect the relative proportions of each lineage (see key). The number of isolates per subregion are denoted near points. (B) Diversity as a function of distance for all isolates and isolates belonging to particular lineages. Individual lineages in each UN subregion were treated as a population and nucleotide diversity (π) was compared to the mean distance of said isolates from the region to Addis Ababa (see *Methods*); population groupings resulting in less than seven isolates were not included. Point size reflects the number of isolates (see key). Data points are in Dataset S1.

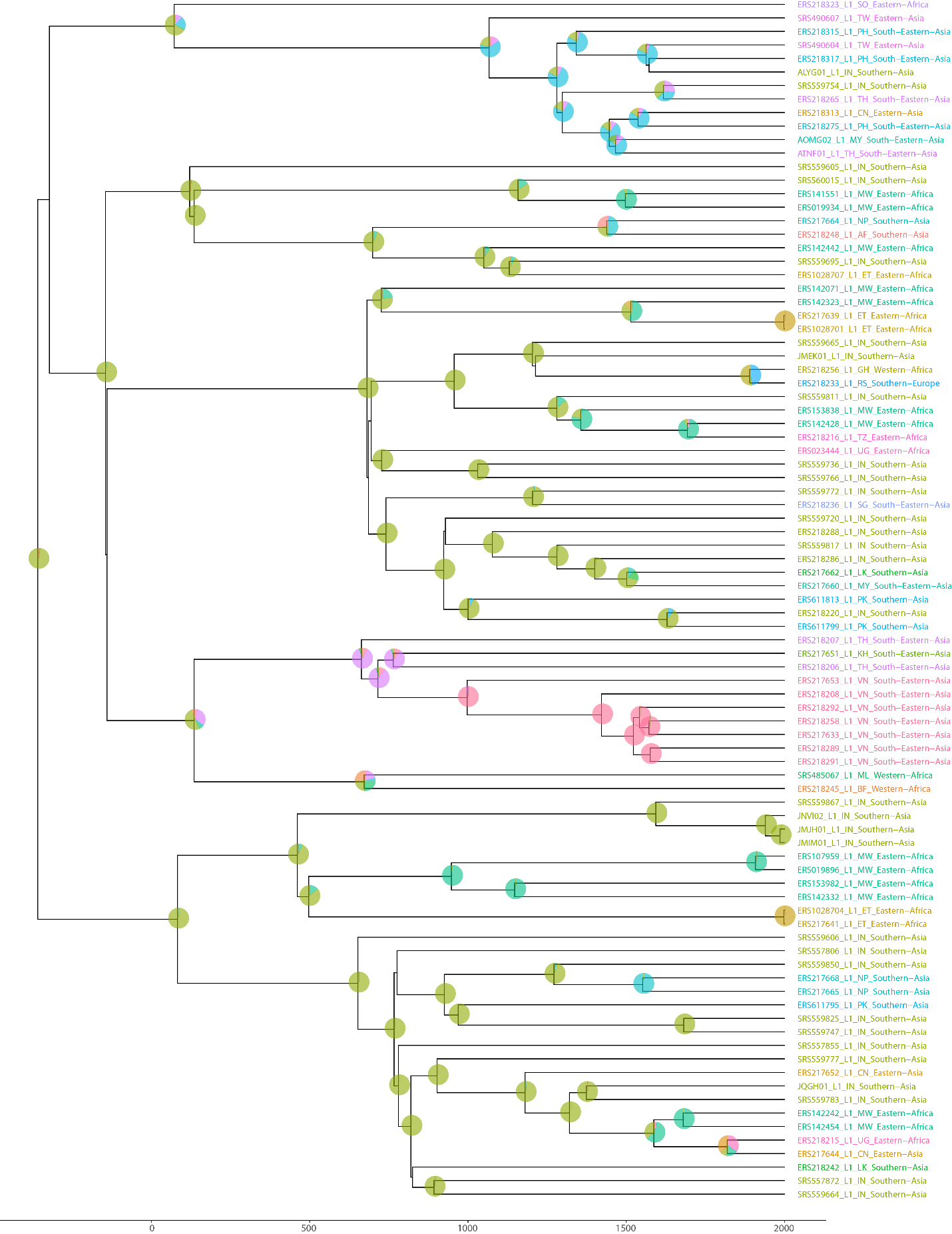

Fig. S7. Maximum clade credibility tree of lineage 1 M.tb genomes. Pie charts at nodes are colored according to the location state probabilities (country). Tip labels are colored according to the country of origin where the isolate was obtained.

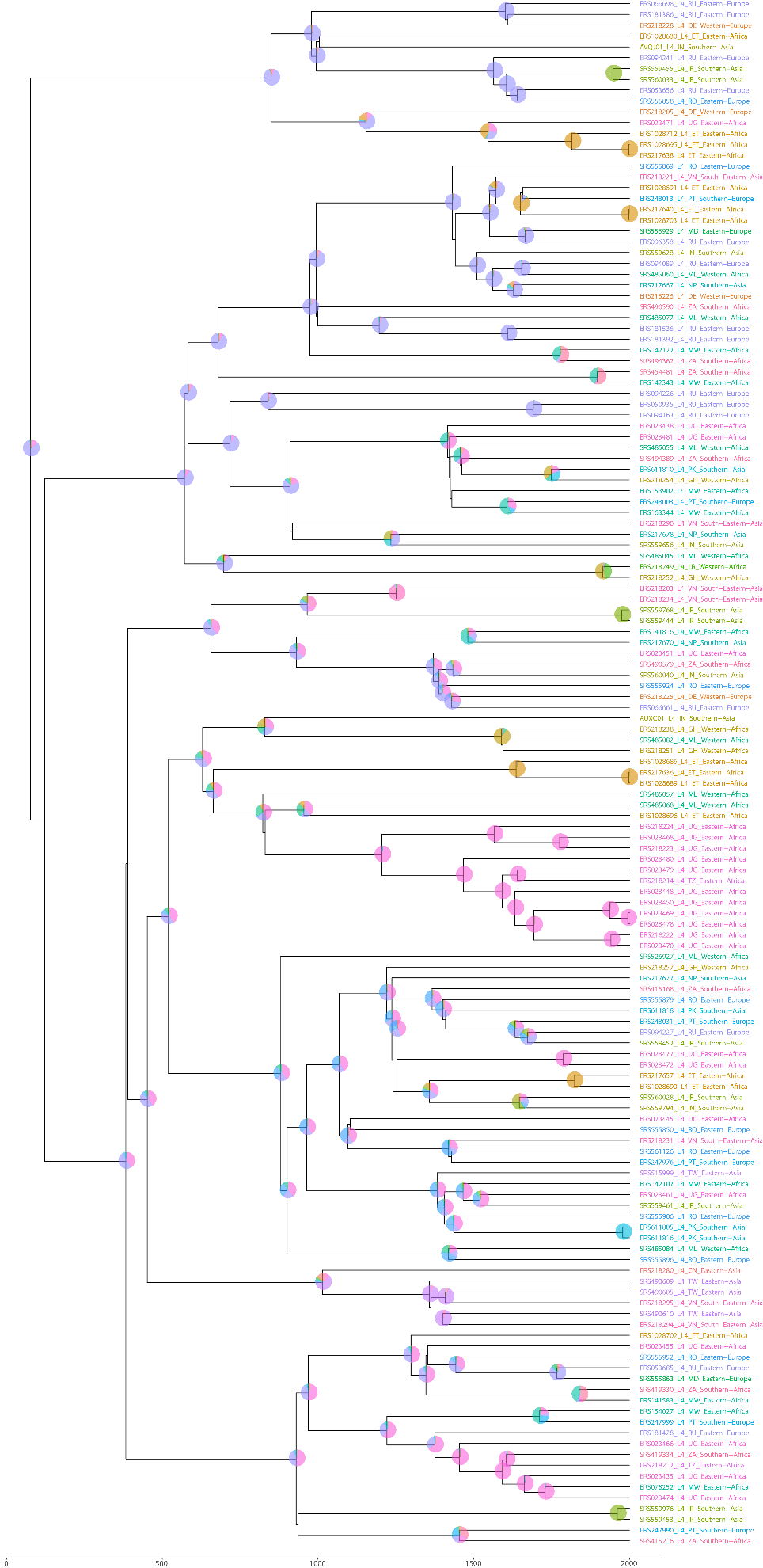
Fig. S8. Maximum clade credibility tree of lineage 4 M.tb genomes. Pie charts at nodes are colored according to the location state probabilities (country). Tip labels are colored according to the country of origin where the isolate was obtained.

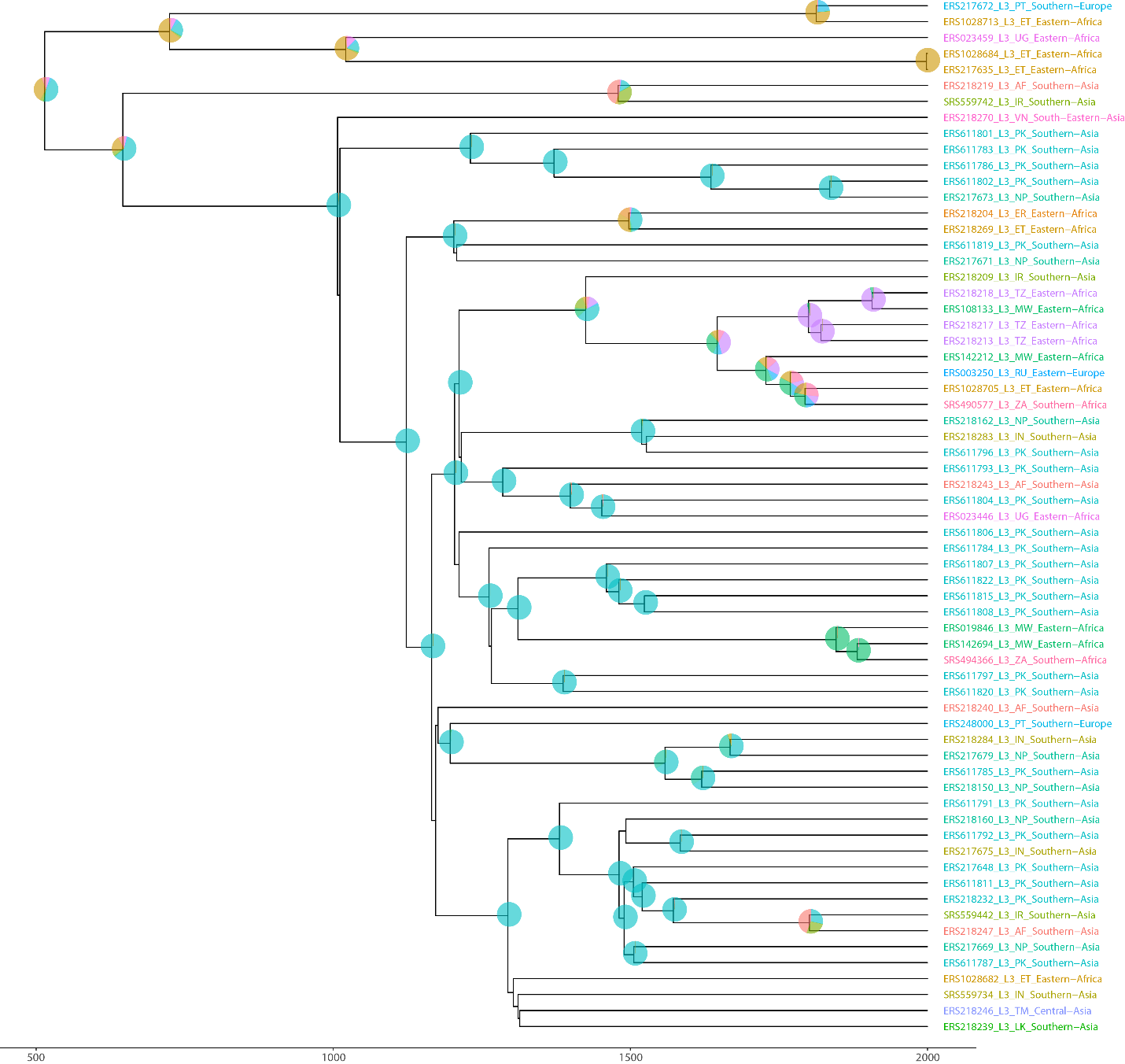

Fig. S9. Maximum clade credibility tree of lineage 3 M.tb genomes. Pie charts at nodes are colored according to the location state probabilities (country). Tip labels are colored according to the country of origin where the isolate was obtained.

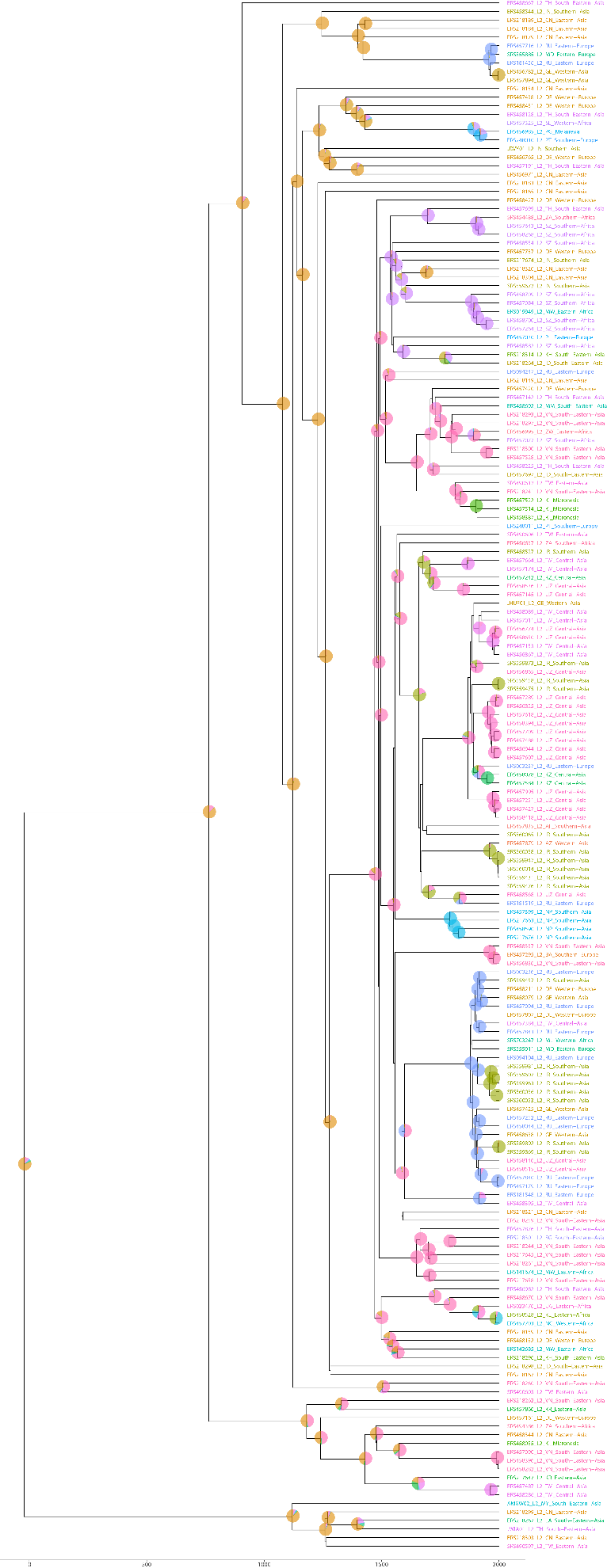
Fig. S10. Maximum clade credibility tree of lineage 2 M.tb genomes. Pie charts at nodes are colored according to the location state probabilities (country). Tip labels are colored according to the country of origin where the isolate was obtained.

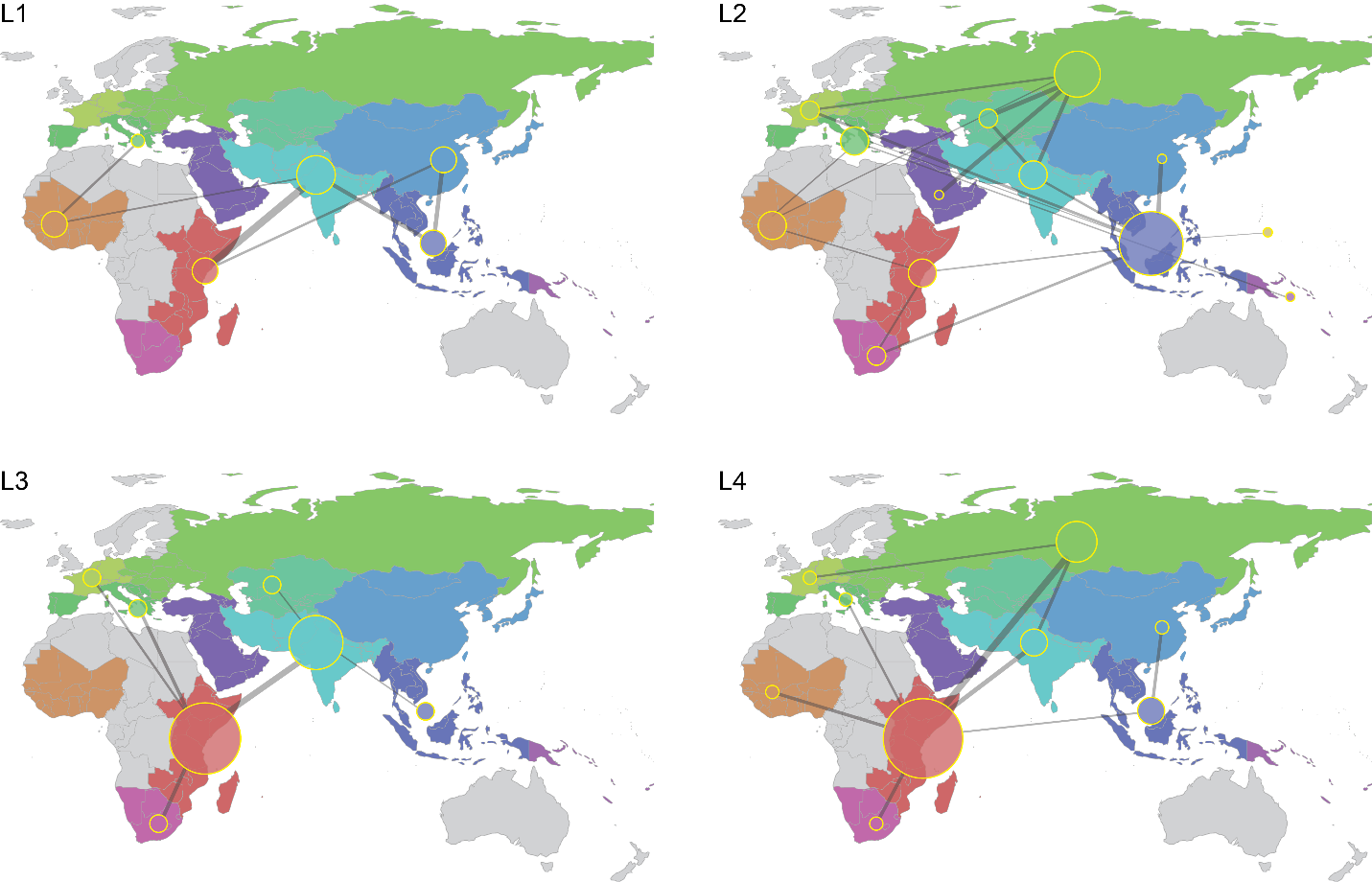

Fig. S11. Relative rates of migration between UN subregions. Relative median rates of migration between geographic regions sampled every 10,000 states in the Bayesian analysis for individual lineages 1-4 are displayed by the thickness of the line connecting the regions. Node size reflects the relative degree of connectivity within each analysis (how many connections the region shares).

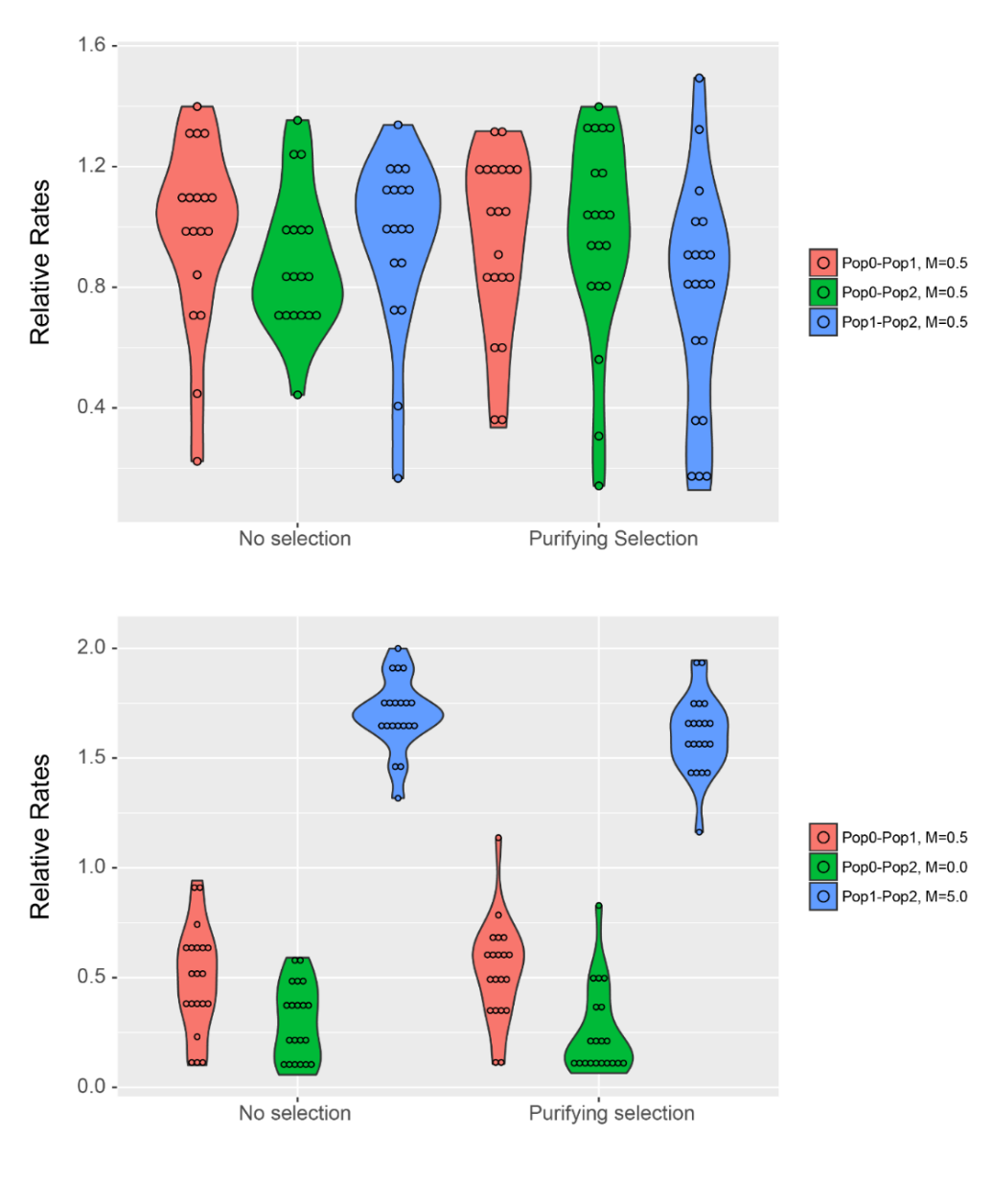

B

A

Fig. S12. Three Population Migration Simulation Rate Estimates. Sequence alignments were simulated using the forward-in-time simulation package SFS_CODE (Hernandez 2008) and analyzed in BEAST (Drummond & Rambaut 2007). Alignments were simulated under (A) symmetric and (B) asymmetric migration regimes, as well as with and without purifying selection. The three possible pairwise migration rates (between populations 0 and 1 = Pop0-Pop1, between 0 and 2 = Pop0-Pop2, and between 1 and 2 = Pop1-Pop2) were all set to 0.5 (i.e., *M* = 0.5) in the symmetric migration regime. In the asymmetric regime, the alignments were simulated with different pairwise population migration rates (Pop0-Pop1 = 0.5, Pop0-Pop2 = 0.0, and Pop1-Pop2 = 5.0). Twenty alignments were simulated with and without purifying selection under each migration regime (total simulated alignments with migration = 80); twenty alignments were also simulated under a scenario without pairwise migration (Pop0-Pop1 = Pop0-Pop2 = Pop1-Pop2 = 0.0) and without purifying selection. The distributions of the migration rate estimates from BEAST analysis are presented as violin plots for the symmetric migration simulations in the top plot and for the asymmetric migration simulations in the bottom plot. There were no statistically significant differences between migration rates estimated from sequences simulated with and without purifying selection.

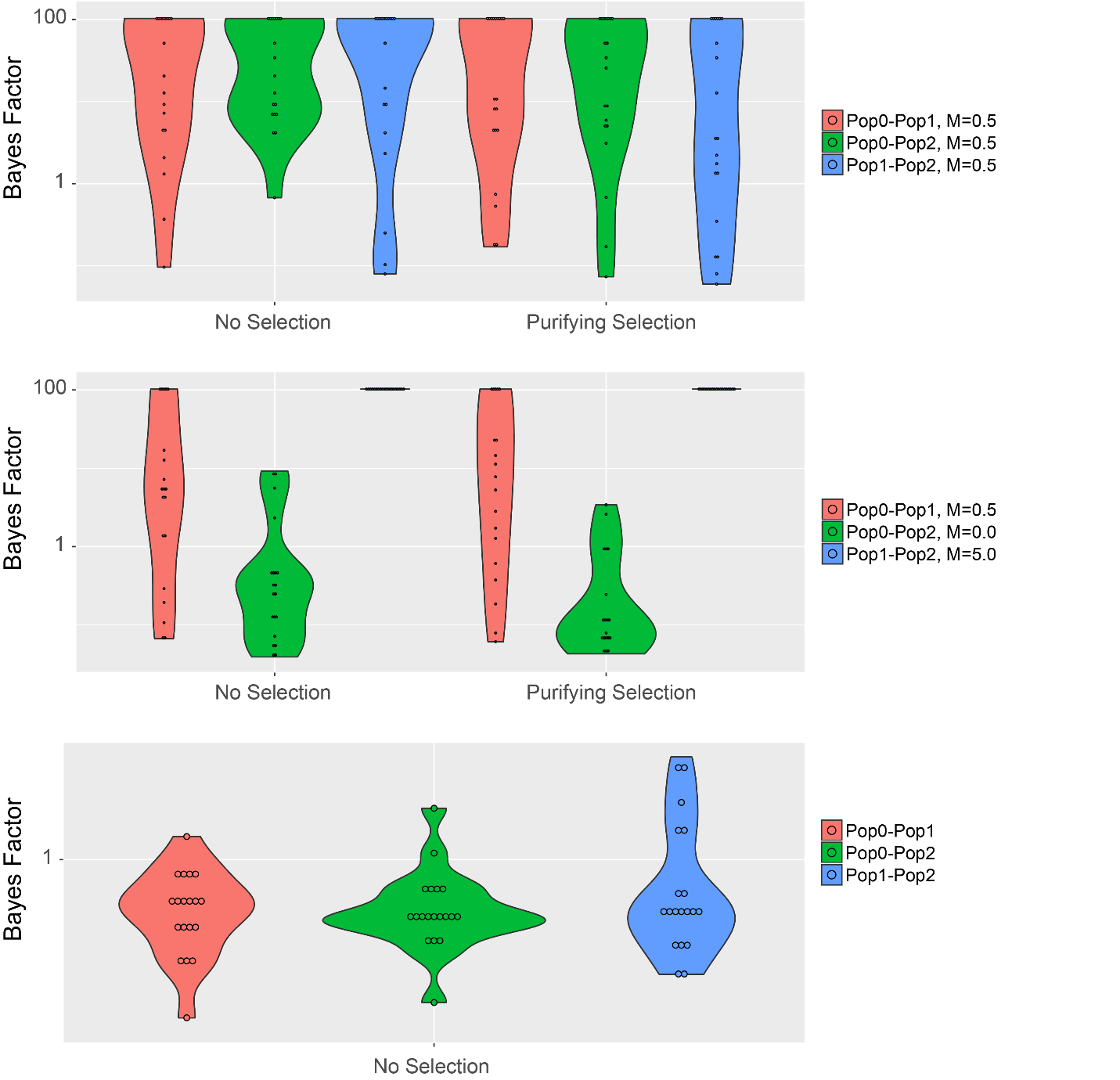

A

B

C

Fig. S13. Three Population Migration Simulation Rate Bayes Factors. Sequence alignments were simulated using the forward-in-time simulation package SFS_CODE (Hernandez 2008) and analyzed in BEAST (Drummond & Rambaut 2007). Alignments were simulated under (A) symmetric and (B) asymmetric migration regimes, as well as with and without purifying selection. The three possible pairwise migration rates (between populations 0 and 1 = Pop0-Pop1, between 0 and 2 = Pop0-Pop2, and between 1 and 2 = Pop1-Pop2) were all set to 0.5 (i.e., *M* = 0.5) in the symmetric migration regime. In the asymmetric regime, the alignments were simulated with different pairwise population migration rates (Pop0-Pop1 = 0.5, Pop0-Pop2 = 0.0, and Pop1-Pop2 = 5.0). Twenty alignments were simulated with and without purifying selection under each migration regime (total simulated alignments with migration = 80); (C) twenty alignments were also simulated under a scenario without pairwise migration (Pop0-Pop1 = Pop0-Pop2 = Pop1-Pop2 = 0.0) and without purifying selection. Bayes factor support for each pairwise population migration rate (i.e., Pop0-Pop1, Pop0-Pop2, and Pop1-Pop2 following the naming convention described in the legend) was estimated using BSSVS (Lemey et al. 2009) in the SpreaD3 package (Bielejec et al. 2016).

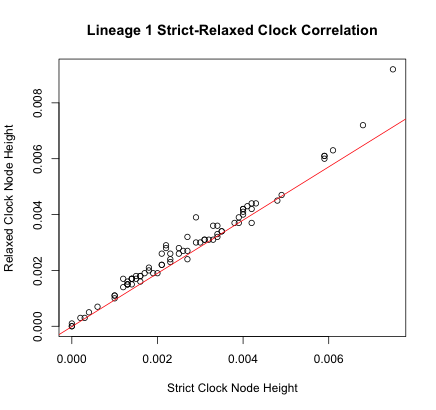

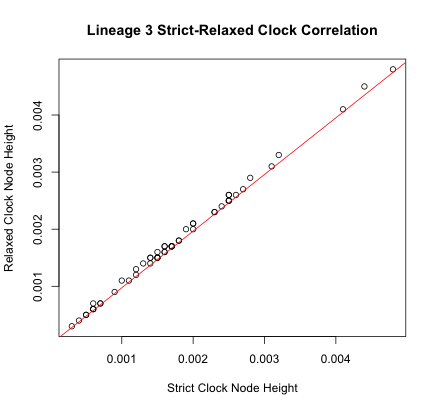

B

A

Fig. S14. Comparison of strict and relaxed clocks. (A) Correlation of Lineage 1 node heights estimated under strict and relaxed (uncorrelated lognormal clock). R^2^ for the correlation is 0.9534. Analyses were performed in BEAST using the same conditions as described in the main text. (B) Correlation of Lineage 3 node heights estimated under strict and relaxed (uncorrelated lognormal clock). R^2^ for the correlation is 0.9925. Analyses were performed in BEAST using the same conditions as described in the main text.

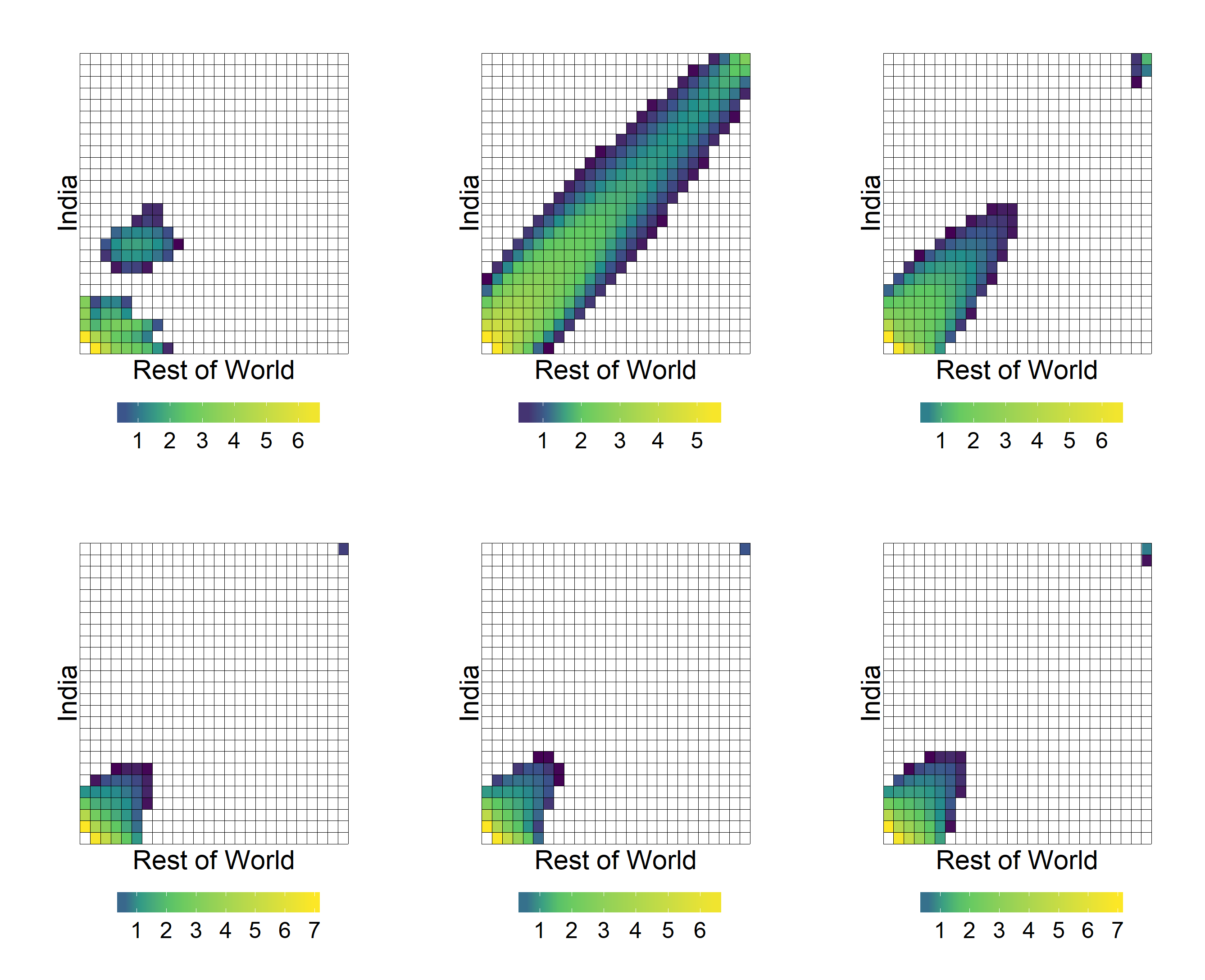

Fig. S15. Observed and inferred 2D SFS of India and the rest of the world (RoW). Heatmaps of 2D spectra colored by number of SNPs at each frequency in the population (log-transformed). SNP counts = 0 in white. From left to right, top to bottom: observed, no split and no migration, no migration, symmetric migration, unidirectional migration, and asymmetric migration.

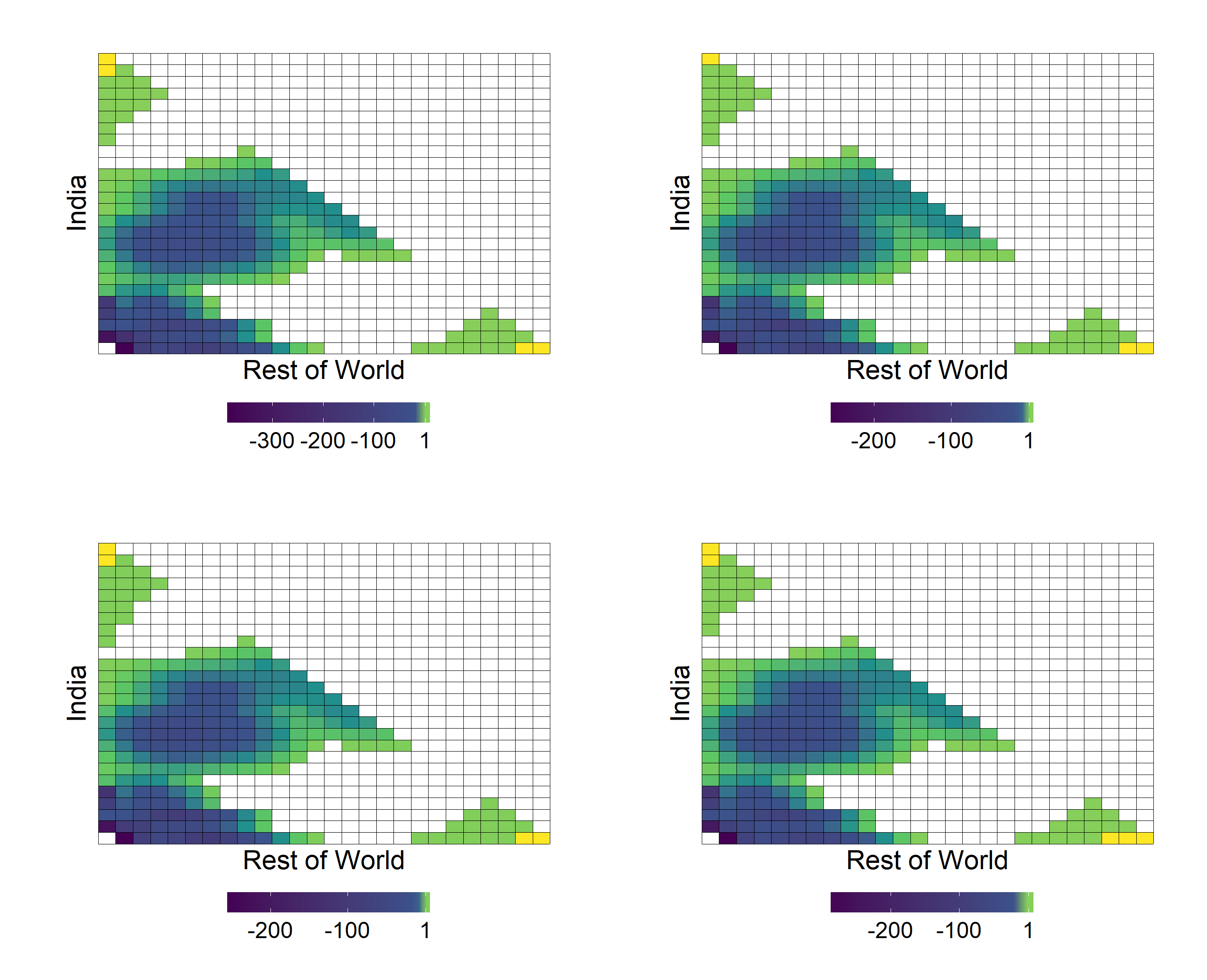

Fig. S16. Poisson residuals for best-fit migration models. Heatmaps of Poisson residuals between model and data for the 2D spectra calculated using ∂a∂I. Masked data colored in white. Positive or negative residuals indicate the model predicts too many or too few SNPs at a given frequency, respectively. From left to right, top to bottom: no migration, symmetric migration, unidirectional migration, and asymmetric migration.

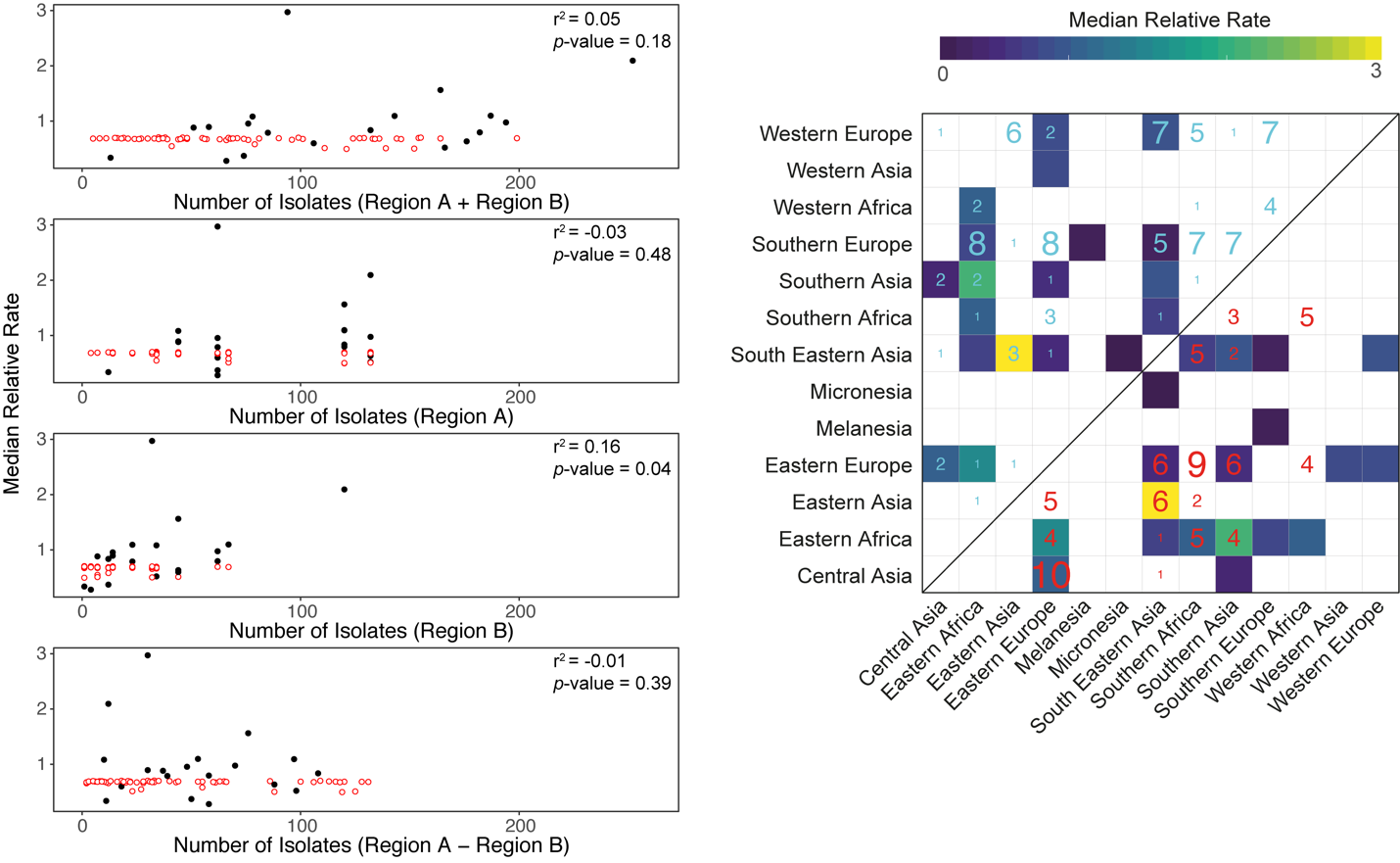

Fig. S17. Relationship between estimated migration rates and sample number. The median relative rate obtained from analysis of the Old World collection with BSSVS (*y*-axis) versus the number of isolates in the Old World collection, where Region A refers to the UN subregion with the most isolates in the comparison and Region B refers to the UN subregion with the least (*x*-axis). Adjusted r^2^ and *p*-values for linear regressions of significant rates (Bayes factor > 5, black circles) are displayed. Open red circles correspond to rates for which there was low support (Bayes factor < 5).

Table S4. Summary of BEAST analyses presented. All analyses were performed using the general time reversible model of nucleotide substitution with a gamma distribution to account for rate heterogeneity between sites and a strict molecular clock. Demographic model, discrete traits, and whether the Bayesian stochastic search variable selection method (BSSVS) was implemented are listed in the table as well as the results derived from the respective analyses.

| **Dataset** | **Demographic Model** | **Discrete Trait** | **Molecular Clock** | **BSSVS** | **Results Derived** |
| --- | --- | --- | --- | --- | --- |
| Old World collection | BSP | UN subregion | strict | Yes | TMRCAs, phylogeography (UN subregion), migration through time, Relative rates of migration between UN subregions |
| Individual lineages (1-7) | BSP | UN subregion | strict | No | Tree structures, BSPs |
| Individual lineages (1&3) | BSP | UN subregion | relaxed | No | Tree structures, BSPs |
| Individual lineages (1-4) | constant | ISO2 code (country) | strict | No | Phylogeography (country) |
| Individual lineages (1-4) | BSP | UN subregion | strict | Yes | Relative rates of migration between UN subregions |
| Two population simulation, 20 replicates of each of 3 scenarios | constant | Population | strict | Yes | Effect of purifying selection on estimates of migration |
| Three population simulation, 20 replicates of each of 5 scenarios | constant | Population | Strict | Yes | Effect of purifying selection on estimates of migration |

Table S5. Summary of ∂a∂i migration analyses for lineage 1. Values for migration model parameters N_e_/N_anc_ (ν), migration (m) and generations since population split (τ), and log_10_ likelihood values (LL) and AIC of each migration model

| Model | v_India_ | v_RoW_ | m_IndiaToRoW_ | m_RoWToIndia_ | Generations | LL | AIC |
| --- | --- | --- | --- | --- | --- | --- | --- |
| **Neutral** | - | - | - | - | - | -2244 | - |
| **No Migration** | 101 | 30 | 0.00 | 0.00 | 0.65 | -507 | 1020 |
| **Symmetric Migration** | 101 | 30 | 0.33 | 0.33 | 1.93 | -477 | 1017 |
| **Unidirectional Migration** | 101 | 30 | 0.72 | 0.00 | 1.68 | -503 | 963 |
| **Asymmetric Migration** | 101 | 30 | 0.01 | 0.65 | 1.56 | -465 | 940 |

Table S6. Summary of ∂a∂i migration analyses of simulated populations. Simulation conditions, the proportion of times ∂a∂i correctly inferred migration conditions of simulations, the model most often selected and the best fit model across replicates (*n* = 20), the best fit model’s LL and AIC.

| Simulation Conditions | Proportion of Migration Conditions Correctly Inferred | Model Most Often Selected Across Replicates | Best Fit Model Across Replicates | Best Fit Model LL | Best Fit Model AIC |
| --- | --- | --- | --- | --- | --- |
| **No Migration** | 0.95 | No Migration | No Migration | -341 | 685 |
| **Symmetric Migration** | 0.25 | Asymmetric | No Migration | -368 | 741 |
| **Symmetric Migration with Purifying Selection** | 0.40 | Unidirectional Migration | Unidirectional Migration | -385 | 775 |
| **Symmetric Migration with Asymmetric Purifying Selection** | 0.10 | Unidirectional Migration | No Migration | -454 | 913 |
| **Unidirectional Migration** | 0.00 | Asymmetric Migration | No Migration | -784 | 1575 |
| **Unidirectional Migration With Purifying Selection** | 0.00 | No Migration | No Migration | -276 | 538 |
| **Unidirectional Migration with Asymmetric Purifying Selection** | 0.10 | No Migration | No Migration | -412 | 828 |
