## Supplementary figures and images for "Lineage specific histories of *Mycobacterium tuberculosis* dispersal in Africa and Eurasia"

### fig. S6

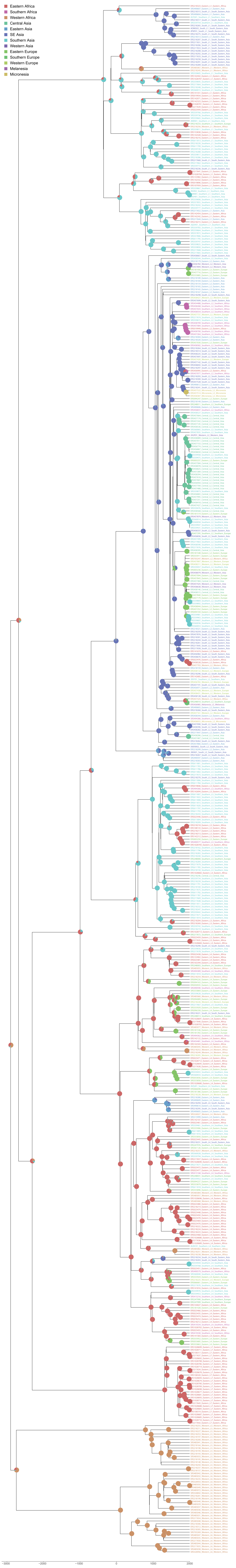
